# Choose carefully, act quickly: Efficient decision making with selective inhibition in attractor neural networks

**DOI:** 10.1101/2021.10.05.463257

**Authors:** Belle Liu, Chung-Chuan Lo, Kuo-An Wu

## Abstract

The ability to decide swiftly and accurately in an urgent scenario is crucial for an organism’s survival. The neural mechanisms underlying the perceptual decision and trade-off between speed and accuracy have been extensively studied in the past few decades. Among several theoretical models, the attractor neural network model has successfully captured both behavioral and neuronal data observed in many decision experiments. However, a recent experimental study revealed additional details that were not considered in the original attractor model. In particular, the study shows that the inhibitory neurons in the posterior parietal cortex of mice are as selective to decision making results as the excitatory neurons, whereas the previous attractor model assumes the inhibitory neurons to be unselective. In this study, we investigate the attractor model with selective inhibition (selective model), which can be considered as a general case of the previous attractor model (unselective model). Our model reproduces several behavioral and neurophysiological observations. To analyze the dynamics of the selective model, we reduce it and show that selectivity adds a time-varying component to the energy landscape. This time dependence of the energy landscape allows the selective model to integrate information carefully in initial stages, then quickly converge to an attractor once the choice is clear. This results in the selective model having a more efficient speed-accuracy trade-off that is unreachable by unselective models.

## Introduction

Decision making is a central cognitive function for animals, and the ability to integrate information in a swift and accurate manner is of great importance. The decision behavior is typically measured by speed and accuracy of the subject, and several computational or mathematical models have been proposed to account for these two quantities (Wang, 2012, 2008; Rao, 2010; Beck et al., 2008; Machens et al., 2005; Lo and Wang, 2006; Bogacz et al., 2006; Fudenberg et al., 2020). In addition to the psychometric measures, the neurological basis of decision making has always been a highly sought-out topic of interest. The attractor neural network models (Wang, 2002; Wong et al., 2007; Wong and Huk, 2008) are the models of interest because they provides insight into the neuronal interactions during the decision process. Furthermore, they link the mathematical equations and variables to known biological substrates, which is crucial for studying how structural changes of the network influences its functionalities.

The attractor model is composed of two competing excitatory populations and one inhibitory population, and all populations are interconnected with one another (Fig. 1a, rightmost). The two excitatory populations receive input signals, each representing “evidence” in favor of one of the two choices, and eventually one population “wins” by silencing the other (Fig. 1b). Attractor models have been successful in recreating the decision making process observed in experiments. One commonly used experimental task is the random dot motion coherence test, where several dots randomly drift towards the left or right (Pilly and Seitz, 2009; Mazurek et al., 2003; Huk and Shadlen, 2005). The subject, upon observing the motion, has to report the direction in which the majority of dots are moving towards.

**Figure 1:**
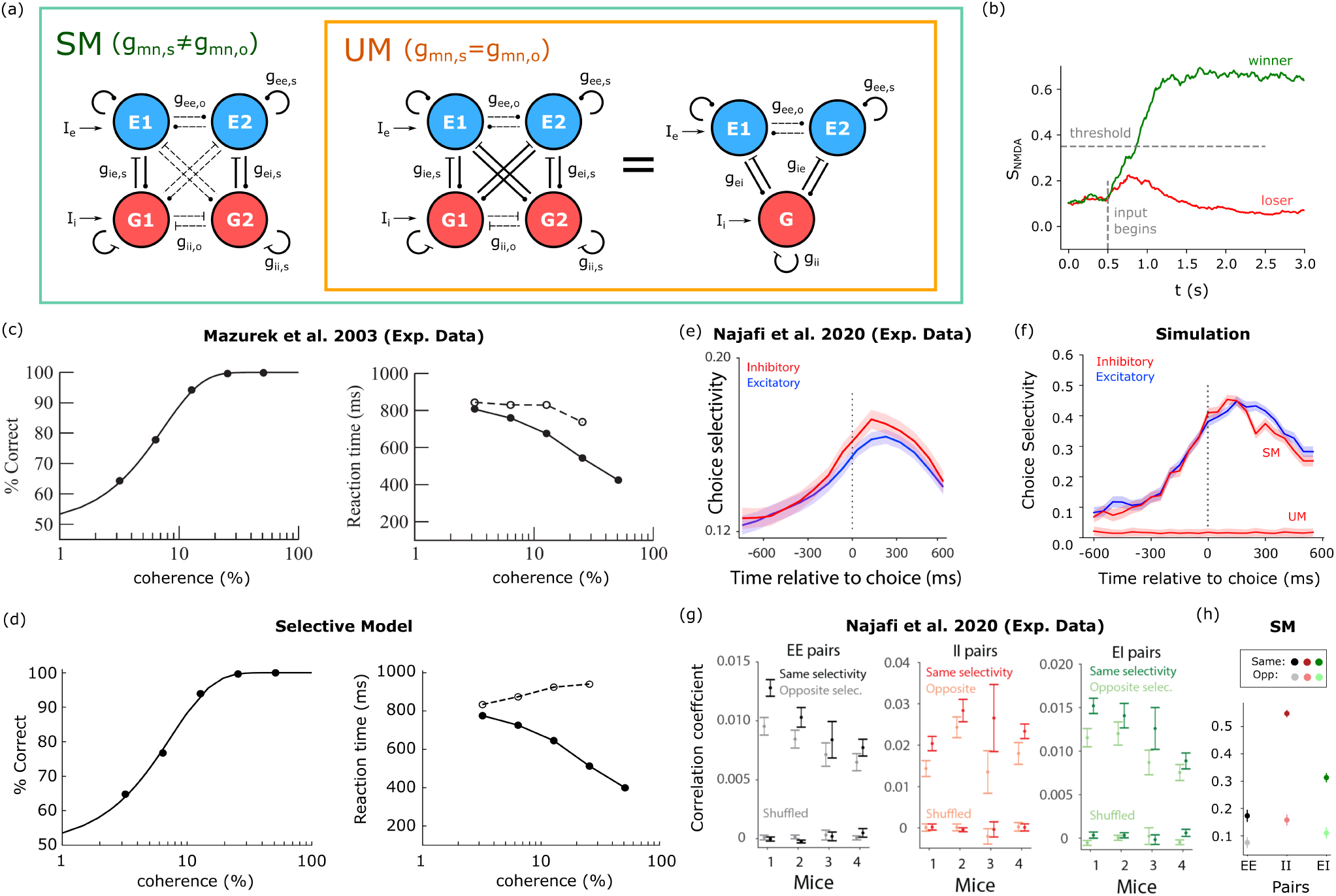
The selective model of decision making can qualitatively reproduce behavioral and physiological observations. (a) The selective model (SM) is a generalized attractor model, with the previous unselective model (UM) being a special case where all inhibitory neurons are homogeneous in synaptic weights. From *left* to *right*: SM, SM with excitatory-to-inhibitory and inhibitory-to-excitatory connections being identical across the two inhibitory populations, which is equivalent to the UM. The use of solid and dashed lines is to highlight the difference in synaptic weights between the populations. (b) Example simulation trial of the two-variable reduced model. The green line represents the activity of the winning population, and red the losing one. The decision is defined as one of the populations’ activity reaching a certain threshold. (c) Percentage correct and mean reaction time for experimental data (Mazurek et al., 2003), figure adapted from (Wong et al., 2007) with permission (Copyright [2006] Society for Neuroscience). (d) Same as in (c) but for SM with {*γ*_*ee*_, *γ*_*ei*_, *γ*_*ie*_, *γ*_*ii*_} = {0.5, 0.6, 0.7, 1}. The *α*_*p*_, *β*_*p*_ values are 7.4 and 1.31 respectively, compared to the experimental values of 7.4 and 1.3. An additional 100 ms is added to the reaction time as the non-decision time that represents other neural processes involved but not related to the decision network modeled here. Number of trials ran is described in Methods. (e) Selectivity as a function of time in experimental data, adapted from (Najafi et al., 2020) with permission. (f) Selectivity as a function of time in the full neural network version of the SM and UM (*γ*_*ee*_ = 0.5). An additional bias is given to the inhibitory populations 150 ms after it has reached its decision, to account for the system resetting itself (see Methods). Choice selectivity is defined as in (e), which is 2 *×* |*AUC* − 0.5| (see (Najafi et al., 2020)); mean *±* SEM by bootstrapping, number of trials *N* = 120. (g) Correlations for EE, II, EI pairs of neurons with either the same (darker color) or opposite (lighter color) selectivity, compared to shuffled data (at the bottom of the graphs). Figure adapted from (Najafi et al., 2020) with permission. (h) Correlations of EE, II, EI pairs of neurons in the full neural network version of the SM. The color schemes mimic those of (g); mean *±* SEM, *N* = 12.

However, new experimental results have revealed new details regarding the network structure of attractor models in the posterior parietal cortex of mice. Specifically, a recent study by Najafi *et al*. (2020) investigated the selectivity of decision making networks (Najafi et al., 2020). Selectivity is important for our understanding of cortical network architecture. Previous studies lean towards a framework where excitatory neurons are selective, but not inhibitory neurons (Sohya et al., 2007; Niell and Stryker, 2008; Kerlin et al., 2010; Bock et al., 2011; Hofer et al., 2011). This is motivated by inhibitory neurons having denser input than excitatory neurons, thereby having broader tuning curves (Hofer et al., 2011). It is also observed that excitatory neurons tend to connect more to other excitatory neurons with similar feature preference, and that its selectivity to orientation increases when the connectivity decreases (Hofer et al., 2011). However, plenty of contradictory results are also reported, where it is found that inhibitory neurons across multiple brain areas respond selectively to different features of the stimulus (Znamenskiy et al., 2018; Runyan et al., 2010; Moore and Wehr, 2013; Allen et al., 2005; Pinto and Dan, 2015; Lovett-Barron et al., 2014; Ego-Stengel and Wilson, 2007).

The aforementioned study by Najafi *et al*. (2020) tested whether selectivity exists within the posterior parietal cortex of mice, using a two-alternative forced choice task (Najafi et al., 2020). Their results showed that the inhibitory neurons are as selective as the excitatory neurons. For the attractor model, the excitatory neurons are selective to the results of the decision. The inhibitory neurons, on the other hand, will react similarly in both cases. Therefore, in the attractor model, excitatory neurons are selective to input while the inhibitory neurons are not. The results of Najafi *et al*. (2020) thus suggests a different network architecture than the previous attractor model, one in which the inhibitory neurons are differently activated by the two inputs.

One of the proposed alternative is a model in which there consists of two inhibitory populations, each receiving stronger input from one population (Fig. 1a, leftmost). This could be thought of as a natural generalization of the attractor model, where inhibitory neurons are allowed to be heterogeneous. The previous attractor model (unselective model, UM) is thereby a special case where all inhibitory neurons are homogeneous in synaptic properties. That is, in the selective model (SM), the synaptic connections from an excitatory population to an inhibitory population (or vice versa) on the same side is not constrained to be the equal to those to the opposite side. In the UM however, they are required to be equal for the inhibitory population to remain unselective, which is an extra restriction.

Our question of interest, following this proposed alternative, is that whether the incorporation of selectivity brings about any computational changes. That is, does the selective model provide computational advantages not present within the unselective model? In this study, we demonstrated how selectivity utilizes a time-varying energy landscape to reach an efficient speed-accuracy trade-off. In particular, the time-varying energy landscape allows the SM to evaluate evidence carefully in initial stages, converge quickly once the choice is clear, and avoid indecisive trials.

## Methods

### Population Models

The classic attractor neural network model of decision making consists of hundreds of neurons, and in combination with the synapses, the complete system will contain thousands of differential equations, which complicates both simulation and analysis (Wang, 2002). In a study by Wong and Wang (Wong and Wang, 2006), it is shown that the attractor model can be reduced to a simpler two-variable form. The reduction takes advantage of several facts: (I) the average activity of the population can be represented by one single variable through mean field reduction; (II) the inhibitory population responds linearly to input; and (III) the time scale of AMPA, GABA and the firing rates are much faster than NMDA, and thus can be assumed to have reached steady state instantaneously. We will call this two-variable model the WWM (Wong and Wang’s Model). The resulting model contains only equations for the NMDA channels of the two competing populations (see Supplementary Notes S1 and Fig. S1). However, since the time constant of noise in the input (Eq S5) is comparable to the time constant of GABA dynamics (2 ms and 5 ms respectively), the rate of change in NMDA could be much shorter compared to its relaxation time. Therefore, the GABA dynamics is retained in the following discussion, and it is shown that including GABA dynamics could alter the dynamics drastically (Supplementary Notes S2, Fig. S2).

Here we will start with deriving the simpler model, which is the UM, and generalize to the SM. With the inclusion of GABA dynamics alongside NMDA dynamics, the full UM is:

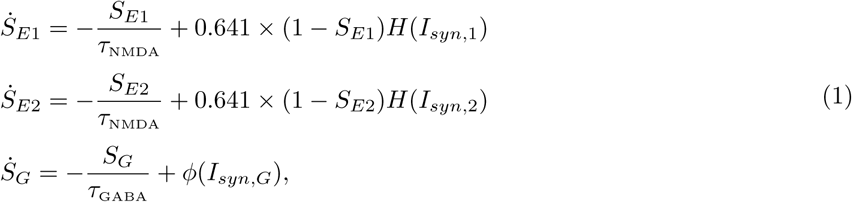

where *S*_*E*1_, *S*_*E*2_ are the gating variables of NMDA, and *τ*NMDA = 0.1 (s) is the time constant of the NMDA channels. Gating variables reflect the degree in which an ion channel is opened. *S*_*G*_ is the gating variable of GABA, and *τ*_GABA_ = 0.005 (s). *H*(*x*) is the *f* − *I* curve of the excitatory population, and is shaped like the lower end of a sigmoid-like function (Supplementary Notes S1). *ϕ*(*x*) takes a form similar to *H*(*x*), but can be linearized since according to experimental findings, the activity of the inhibitory population never falls into the nonlinear regime (i.e. assumption (II)) (Wong and Wang, 2006). The synaptic current, *I*_*syn*_, is a weighted sum of pre-synaptic inputs.

The terminology for the synaptic weights are shown in Fig. 1a. To briefly summarize, for a synaptic weight *g*_*mn*_ where {*m, n*} ∈ {*e, i*}, *m* represents the pre-synaptic population type (*e* for excitatory, *i* for inhibitory) and *n* the post-synaptic population type. The intra-population excitatory-to-excitatory synaptic weight (*g*_*ee,s*_, where *s* stands for *same side*) is different from the inter-population excitatory-to-excitatory synaptic weight (*g*_*ee,o*_, where *o* stands for *opposite side*). The bias currents to the excitatory and inhibitory populations are denoted as *I*_*e*_ and *I*_*i*_.

For simplicity, the inhibitory populations receive no noise. The equations for the UM synaptic currents are:

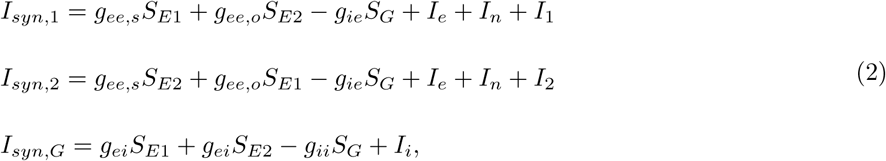

where *I*_1_, *I*_2_ represent the input signal, and *I*_*n*_ is a Gaussian white noise (Supplementary Notes S1).

The SM essentially divides the inhibitory population into two, allowing them to take on different synaptic weights. Therefore, the equation for the SM is:

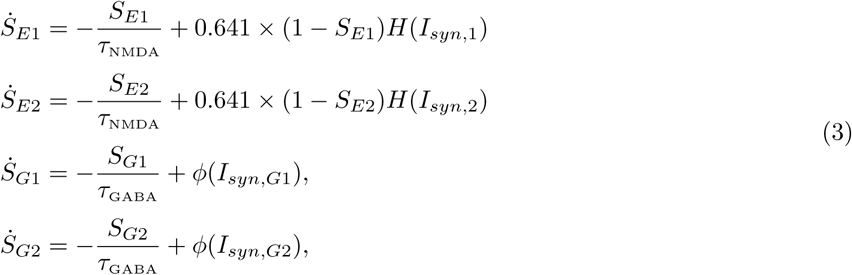

The SM synaptic currents are:

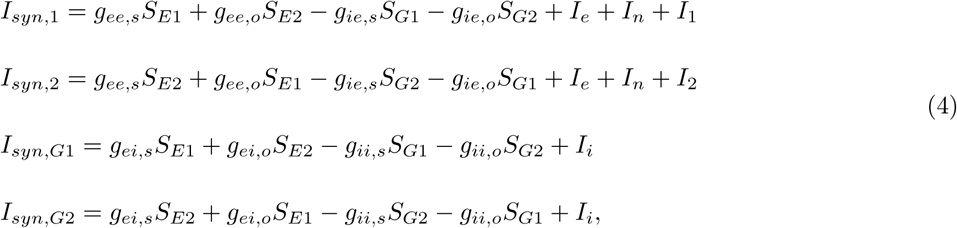

To ensure fair comparison across different models that we study, and to isolate the effect of selective inhibition, we created models whose equilibrium points are the same when projected onto the *S*_*E*1_-*S*_*E*2_ plane. That is, upon activating assumption (III), these model would all collapse back into an identical WWM. This provides an additional benefit of assuring us that GABA dynamics do in fact make a difference; because if it does not, then all models should yield the same results since their two-variable form is identical.

A technical description of how this is done is as follows: for the UM, enforcing assumption (III) on Eq 1 yields

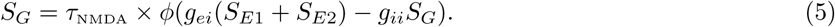

That is, since assumption (III) assumes that *S*_*G*_ always reaches the equilibrium state immediately, it assumes that 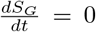, which is how we were able to arrive at Eq 5. By substituting Eq 5 back into Eq 1, we can once again lower the number of differential equations back to two. Since in Eq 5, *S*_*G*_ is a linear combination of *S*_*E*1_ and *S*_*E*2_, after substitution the synaptic currents *I*_*syn*,1_ and *I*_*syn*,2_ will remain a linear combination of *S*_*E*1_ and *S*_*E*2_. Therefore, we could collect the coefficients for *S*_*E*1_, *S*_*E*2_ and the bias term in the synaptic currents – which would consist of the parameters {*g*_*ee,s*_, *g*_*ee,o*_, *g*_*ei*_, *g*_*ie*_, *g*_*ii*_, *I*_*e*_, *I*_*i*_} – and call them {*J*_*e*_, *J*_*i*_, *I*_*b*_}, which are the parameters for the WWM (Fig. S1). *J*_*e*_ represents the strength of self-excitation, *J*_*i*_ represents the degree in which the competing populations inhibit one another, and *I*_*b*_ is the bias.

Since the degrees of freedom in the former exceeds the latter, there are infinite sets of UM parameters {*g*_*ee,s*_, *g*_*ee,o*_, *g*_*ei*_, *g*_*ie*_, *g*_*ii*_, *I*_*e*_, *I*_*i*_} that could be mapped to the same two-variable parameters {*J*_*e*_, *J*_*i*_, *I*_*b*_}. For convenience purposes, four constraints will be added to the mapping process, so that the mapping becomes one-to-one. This allows us to be able to adjust one single parameter, and have the other parameters be automatically sought out. These four constraints are: (I) *g*_*ie*_ = *g*_*ei*_, (II) *g*_*ii*_ = 1.2, (III) *I*_*i*_ = 1, and (IV) 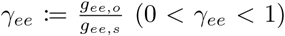. The first constraint originates from the observation that excitation and inhibition are often balanced in neural networks (Najafi et al., 2020; Lam et al., 2017; Wang et al., 2013). The second and third are chosen so that the resulting UM does not violate assumption (II). Constraint (IV) defines a parameter *γ*_*ee*_ for which we can adjust to control the speed in which the activity of the excitatory population ramps up. Note that when *γ*_*ee*_ is altered, the synaptic weights {*g*_*ee,s*_, *g*_*ee,o*_, *g*_*ei*_, *g*_*ie*_}, as well as bias currents {*I*_*e*_, *I*_*i*_} will all change along in order to satisfy the requirement imposed by assumption (III) and the four constraints.

Under these constraints, we can obtain unique UM parameters (Supplementary Notes S2). A reduction from the SM to the WWM can also be conducted in a similar fashion. That is, the differential equations for *S*_*G*1_, *S*_*G*2_ can be simplified in the same manner as Eq 5, and again substituted into Eq 3, which would again lead to a WWM (Supplementary Notes S2).

Similar to how *γ*_*ee*_ was defined in the UM to account for the ratio between the inter- and intra-population synaptic weight of the excitatory-to-excitatory connection, we also define 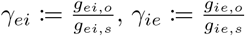 and 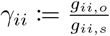 for the other three types of connections that exist in the SM. Note that while *γ*_*ee*_ must be strictly between 0 and 1 in order for the network to function properly, no such constraints exist for *γ*_*ei*_, *γ*_*ie*_ and *γ*_*ii*_. Intuitively, if *γ*_*ee*_ *>* 1, then the favored population would excite its opponent more than it excites itself (*g*_*ee,o*_ *> g*_*ee,s*_), leading to a situation where no decision can be made. However, if say *γ*_*ie*_ *>* 1, it merely means that the inhibitory population prefers to inhibit the contralateral excitatory population more; if *γ*_*ie*_ *<* 1, then the inhibitory population inhibits its ipsilateral excitatory population more.

### Reduced Population Models

The UM and SM have three and four variables respectively, which makes geometrical interpretations of their dynamics difficult. To better analyze the UM and SM, we reduced the models back to a two-variable form. However, instead of using assumption (III), the reduction is based on empirical observations of the simulations of population models, which allows the reduced models to better capture the dynamics of the full models.

#### Linear Relation of Average Excitation and Average Inhibition in UM and SM

It is observed that *S*_*G*_ and the average excitation *S*_+_ in the UM, where 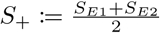, co-evolve in a linear manner throughout the simulation (Fig. S3a, e). An analogous relation arises in the SM as well, where the linear relation is between *S*_*G*_+ (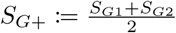, i.e. average inhibition) and *S*_+_. Therefore, they are fitted to a linear function, *S*_*G*_ = *aS*_+_ + *b* (*S*_*G*+_ = *aS*_+_ + *b* for the SM). The goal of fitting a linear function to *S*_*G*_ and *S*_+_ is to reduce the UM and SM back into the two variable form, with new parameters that more accurately describe its dynamics than if assumption (III) were in place. That is, we can replace *S*_*G*_ by *aS*_+_ + *b* in Eq 2, so that the number of variables once again reduces back to two. The synaptic currents after this simplification become:

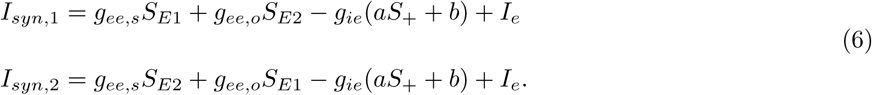

By letting 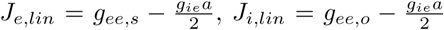 and *I*_*b,lin*_ = *I*_*e*_ − *g*_*ie*_*b*, we note that this model has the exact same mathematical form as WWM (Supplementary Notes S1). We will refer to the parameters of the linear reduced model as {*J*_*e,lin*_, *J*_*i,lin*_, *I*_*b,lin*_}, even though they play the same role as {*J*_*e*_, *J*_*i*_, *I*_*b*_} in the WWM.

The fitted *a, b* values vary slightly across each trial; therefore, a more accurate reduction would be to obtain *a*_*sim*_ and *b*_*sim*_ via averaging over the simulated trials of the UM. However, to avoid having to run numerous trials just to obtain *a* and *b*, they are instead approximated by running a single trial in which no noise is present. The noiseless trial starts at the stable equilibrium point in which *S*_*E*1_ = *S*_*E*2_, and in the absence of noise, the system stays on the separatrix (the line where *S*_*E*1_ = *S*_*E*2_) throughout the simulation. The resulting slope *a*^∗^ and intersect *b*^∗^ is then used to simulate the linear reduced model of the UM.

*a*^∗^, *b*^∗^ obtained by the noiseless trials are compared to *a*_*sim*_, *b*_*sim*_ obtained by simulations to evaluate its validity (Fig. S3b). The data points used for fitting *a*_*sim*_, *b*_*sim*_ are the data points beginning from the onset of the input (*t* = 0.5 (s)) up to *t* = 1.0 (s). This is because including data points where the system is in the stable state only adds noise to the fitting process and no information. Therefore, the time in which the system is in the initial stable state is omitted by only including points from *t* = 0.5 (s) onward. Since the time in which each trial reaches the final stable state fluctuates, we compensated by truncating the data points beyond *t* = 1.0 (s). This is because the average reaction times seldom exceed that value.

#### Quadratic Relation of Excitation Difference and Inhibition Difference in SM

For the SMs, we define excitatory difference 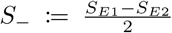 and inhibitory difference 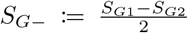. (Note that an analogous term of *S*_*G*−_ does not exist in the UM, since the UM does not contain two inhibitory populations.) While at first glance, the relation between *S*_*G*−_ and *S*_−_ appears linear, it is actually quadratic (Fig. S3c, e). The coefficients of this quadratic relation, 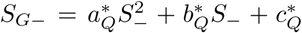, cannot be obtained by running a fair trial (i.e. *S*_*E*1_ and *S*_*E*2_ starting out with the same value) without noise, since this would result in having *S*_−_ = 0 at all times. Instead, an unfair trial without noise is run by having the initial *S*_*E*_ values be biased towards one side ((*S*_*E*1_, *S*_*E*2_) = (0.26, 0.25) at *t* = 0). The data points used to find *a*_*Q,sim*_, *b*_*Q,sim*_ are similarly the ones from the onset up to 1.0 (s), and they are also compared to 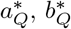 in Fig. S3d. The fact that this quadratic relation exists in a noiseless trial shows that this is not a noise-induced artifact, but a property inherent to the SM.

For analysis purposes, we model this quadratic relation as a linear relation with a time-varying slope and intercept. That is, we linearized the quadratic relation within short periods of time. In the noiseless trial, the data points for every 250 ms are fitted to a line,

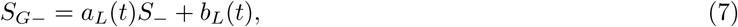

where the coefficients *a*_*L*_(*t*), *b*_*L*_(*t*) change for the different periods of time. Substituting Eq 7 into Eq 4, the synaptic currents become:

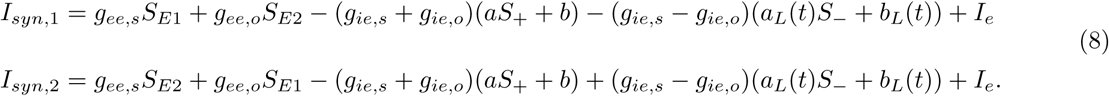

We call this the time-dependent linear reduced model. The only difference this model has with the WWM or the linear reduced model in terms of mathematical form is the time dependence of its variables *a*_*L*_(*t*) and *b*_*L*_(*t*). The relations between the WWM, the full UM/SMs, the linear reduced model and the time-dependent linear reduced model are summarized in Fig. S4. The performance of the reduced models in comparison to the full models are summarized in Supplementary Notes S3 and Fig. S3.

### Full Scale Neural Network Model

Population models, being a simplification, may not be able to capture certain quantities observed in experiments in which activities of individual neurons were measured. Therefore, in order for us to compare the UM and SMs to experimental data, we constructed a full neural network model that contains 50 neurons in each population. The connections are all-to-all between populations or within a population, and the synaptic weights are homogeneous for the same type of connections (e.g. *g*_*ee,s*_ is the same for all excitatory connections that start and end at the same population). The equations of each neuron are the same as the mean field equations (Eq 1, 2), but with each neuron receiving independent noise. The parameters of the model are (*γ*_*ee*_, *γ*_*ei*_, *γ*_*ie*_, *γ*_*ii*_) = (0.5, 0.6, 0.7, 1).

For tuning the full neural network to have similar selectivity trends as the experimental results of (Najafi et al., 2020), we set the input magnitude *µ*_0_ to 0.0052, the excitatory noise standard deviation *D* to 5.0 and the inhibitory noise standard deviation is *D* = 2.5 (see Eq S5). To account for additional events that might happen after the decision, we added a mechanism that resets the network. 150 ms after the network has reached its decision, the excitatory neurons stop receiving inputs, and the inhibitory neurons begin receiving a small bias current of 0.002. This would cause a decline in the network activity, and thus reset the network.

When tuning the network to have similar correlation trends as those of (Najafi et al., 2020), we set the excitatory noise standard deviation *D* to be 6.0, and the inhibitory noise standard deviation is *D* = 1.0. We set a weak correlation coefficient to be 0.05 between the noise inputs of every pairs of neurons in the network. No input aside from noise is given to the network, so as to mimic spontaneous activities. Note that the full neural network is used only to compare to the experimental data of Najafi et al., and for all other experimental comparisons and mathematical analyses, we used the aforementioned reduced models of UM and SM.

### Energy Landscape

To gain a more intuitive understanding of the underlying dynamical system, we calculated an energy-like function *U* (Lyapunov function), which governs the dynamics of gating variables. In particular, the “valleys” of the energy landscape correspond to the stable attractors of the dynamical system, and the “hilltop” the unstable attractors. The dynamics of the system would seek to minimize its value, which means that the system behaves like a ball that “runs towards” the valleys.

In terms of our models, the energy landscape can be solved semi-analytically for the WWM and linear reduced model. The time-dependent linear reduced model, with its richer dynamics, on the other hand exhibits a time-varying energy landscape. Hence, instead of directly calculating *U* in these systems, the energy landscape of all the models will instead be approximated by averaging over simulation trials (Wong and Wang, 2006), and used as a qualitative visualization tool for understanding the properties of the dynamical system.

For a dynamical system 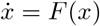, its energy landscape *U* (*x*) is associated to *F* (*x*) by *F* (*x*) = −*dU/dx* (Wong and Wang, 2006; Roxin and Ledberg, 2008). Letting *x* = *S*_*E*1_ − *S*_*E*2_, the value of *F* (*x*) and consequentially the value of *U* (*x*) can be obtained via simulation. *F* (*x*) is simply (*x*(*t* + Δ) − *x*(*t*))*/*Δ, where Δ is the simulation time step. Since the individual trials are noisy, these values are averaged across 2000 trials to obtain a smooth function,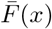. *U*(*x*) is then an integration across 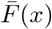 in terms of *x*. The energy landscape obtained would resemble a double-well potential, where the two “valleys” correspond to the two choices presented to the system. The system will be forced to move towards one of the valleys, which corresponds to it making a decision (Fig. 1b).

### Simulation Protocol

The simulations, unless stated otherwise, always start at the stable equilibrium point where *S*_*E*1_ = *S*_*E*2_. The input signals for the two excitatory populations are

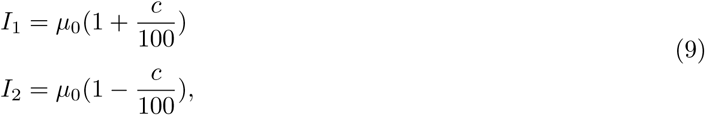

where *µ*_0_ = 0.0156 (nA), and *c* represents the coherence of the signal. A large *c* corresponds to a random dots motion test in which the majority of points move in the same direction (the one that favors population 1). The values of *c* used here are the same as those used in (Lo, 2010): 0, 3.2, 6.4, 12.8, 25.6 and 51.2%. The lower the coherence, the higher the variance of the accuracy. Therefore, the number of trials ran *N* decreases for higher *c*: 1500, 1500, 1500, 1000, 250, 100 are run for the respective coherence levels.

At *t* = 0.5 (s), the input signals are turned on, and the network begins to make a decision. A decision is defined as one of the populations’ NMDA gating variable exceeding a threshold of 0.35 (unless stated otherwise), and the winner is defined as the first population to do so. The reaction time is then the time it takes for the winner to reach the threshold. The total simulation time of a trial is 3 (s), and if the network is undecided until that point, the winner is randomly assigned and the reaction time is marked as 2.5 (s). These undecided events seldom occur, especially when the threshold is low. For reference, none of the UMs had undecided trials out of a total of 1500 trials when the threshold is set to 0.35. When the threshold is increased to 0.6 (the largest threshold used in this study), excluding these undecided trials caused a change of 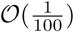 (s) in the reaction time of incorrect trials, but did not make a difference in the reaction time of correct trials.

All of the ODEs are solved by the fourth-order Runge-Kutta method with a time step of 0.1 ms.

### Statistical Analysis

#### Speed-Accuracy Trade-off

A trade-off between accuracy and reaction time seems inevitable, but not all models are efficient in its trade-off. To quantify whether the trade-off of a model is efficient compared to others, we first parameterized the psychometric function by this form (Lam et al., 2017):

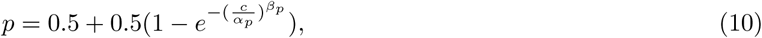

where *p* is the accuracy, *c* the coherence, *α*_*p*_ the discrimination threshold and *β*_*p*_ the psychometric order. Essentially, *α*_*p*_ controls the coherence that hits 81.6% correct, and *β*_*p*_ controls the slope at said threshold. We can then use *α*_*p*_ (we will call it the performance index) as a measurement of accuracy for the model: the smaller it is, the better.

The reaction time, since it decreases monotonically as a function of coherence, will be represented by the average reaction time when coherence equals zero (speed index, *α*_*RT*_). Here, both accuracy and reaction time are bootstrap sampled 1000 times to obtain its standard error.

Finally, we can plot the accuracy versus reaction time for each parameter set. Any parameter set that is on the upper right corner of another parameter set is less efficient. That is, if we have *α*_*RT*_ as the x-axis and *α*_*p*_ as the y-axis, and if point B is located in the upper-right direction of point A, this indicates that it has a longer reaction time compared to parameter set A, yet performs worse.

#### Non-Regret Choice

During the initial stage, the system walks the separatrix, alternating between the basins of attraction of *S*_*E*1_ and *S*_*E*2_. For a given deviation from the separatrix, e.g. *S*_*E*1_ − *S*_*E*2_ = 0.04, the system might continue falling into the favored attractor, *S*_*E*1_, or it might end up in the other one. Conceivably, the larger the deviation, the less likely it is for the system to “change its mind”. We call the value of the deviation a benchmark, in order to distinguish it from the threshold (the threshold is *S*_*E*1_ or *S*_*E*2_ reaching some assigned value; a benchmark is |*S*_*E*1_ − *S*_*E*2_| reaching some assigned value).

For each benchmark, we can calculate the probability of the system ending up in the attractor it currently favors. We call this the non-regret choice percentage. As a trivial example, consider having coherence *c* = 0 and benchmark equal to zero: the non-regret choice percentage is roughly 0.5, because only half of the trials that start out favoring one side will actually converge to that side. Comparing how the non-regret choice percentage changes as a function of the benchmark would reveal how careful the system is weighing its options. For instance, if the percentage is high for a small benchmark value, this means that as soon as |*S*_*E*1_ − *S*_*E*2_| exceeds that benchmark, the system seldom changes its choice.

## Results

### Selective Model Conforms with Experimental Data

To show that the selective model as a generalization of attractor model is more biologically realistic, we compared the SM to some of the existing experimental data. We first look at the basics of a decision making task, which are the psychometric and chronometric functions, by comparing the selective population model to experimental data (Fig. 1c, d). A decision making model can be fitted to random dots motion task data (Roitman and Shadlen, 2002) by adjusting the standard deviation of the noise, *D* (see Eq S5). Both the UM and SM can be reasonably fitted to the experimental data, but the SM requires lower *D* values to achieve the experimental values of *α*_*p*_ and *β*_*p*_, which already hints at the efficiency of the SM.

Aside from the psychometric and chronometric functions, the other important property for a decision making network its selectivity, as shown by Najafi *et al*. (2020). To better fit our selective model to the experimental result, we constructed a full-scale neural network model that contains 50 neurons in each population. The connections are all-to-all, the synaptic weights are identical and homogeneous between populations and within a population, and each neuron receives independent noise. We discovered that the full-scale neural network model exhibits a similar selectivity trend as those seen in the study of Najafi *et al*. (2020) (Fig. 1e, f).

A further prediction of having a SM configuration is that neuron pairs with the same selectivity would have a higher correlation than pairs with opposite selectivity, since their connection weights are stronger. This is shown experimentally by Najafi *et al*. (2020), where correlation of the same selectivity pairs are larger than the opposite pairs, and both are non-negative (Fig. 1g). However, in the simulations, we note that if each neuron received uncorrelated noise, the Pearson correlation of neurons with the opposite selectivity would be negative. This is because in the SM configuration, the two excitatory-inhibitory pairs are naturally in competition with one another, which would result in their activities moving in opposite trends. This argues for the presence of weakly correlated noise between neurons. Indeed, when adding weakly correlated noise, the SM shows the same trend in correlations as the experimental data (Fig. 1h).

An interesting detail to note here is that both in tuning for selectivity and correlation, the inhibitory neurons must receive less noise than the excitatory ones to produce results that are qualitatively similar to the experimental data. For example, if they are given equal amounts of noise, the excitatory-excitatory pairs would end up having a much higher Pearson correlation than the inhibitory-inhibitory pairs, while the opposite is shown in experimental data (Fig. 1e). This is because excitatory neurons tend to synchronize with one another, while inhibitory neurons tend to move in an asynchronous manner.

### Selectivity Improves Speed-Accuracy Trade-Off

While SMs explain experimental data better than the UM, whether selectivity is valuable in a computational sense is still undetermined. To this end, we investigated speed-accuracy trade-off, which is a commonly used metric. An efficient trade-off of speed and accuracy, where the reaction time is low but the accuracy is high, is desirable. Specifically, we tested how speed-accuracy trade-off is influenced by selectivity, which is quantified by the parameters *γ*_*ei*_ and *γ*_*ie*_. Smaller *γ*_*ei*_ and *γ*_*ie*_ values would lead to a larger difference in the inter-population excitation and inter-population inhibition values, and thus to a more selective model. To compare the speed-accuracy trade-off of SM with UMs, we parameterized the psychometric and chronometric functions, and used the performance index *α*_*p*_ to represent accuracy and the speed index *α*_*RT*_ to represent the speed (see Methods and Fig. 2a, b). The smaller the performance index is, the more accurate the model; the smaller speed index is, the faster the model. Therefore, a smaller *α*_*p*_ × *α*_*RT*_ value would indicate a more efficient model, and thus we define the trade-off performance of a model by the inverse of *α*_*p*_ × *α*_*RT*_. Another metric to compare to would be the reward rate, which is simply the total number of rewards across all trials over the total reaction time (in units of *s*). The reward is 1 for each correct trial, and the trials consist of randomly selected coherence levels 0, 3.2, 6.4, 12.8, 32.6, 51.2, where each coherence level is equally likely to be selected. For *c* = 0 trials, 50% of the trials are randomly assigned as correct. The trend of efficiency of the trade-off are similar across the two metrics (Fig. 2c, d).

**Figure 2:**
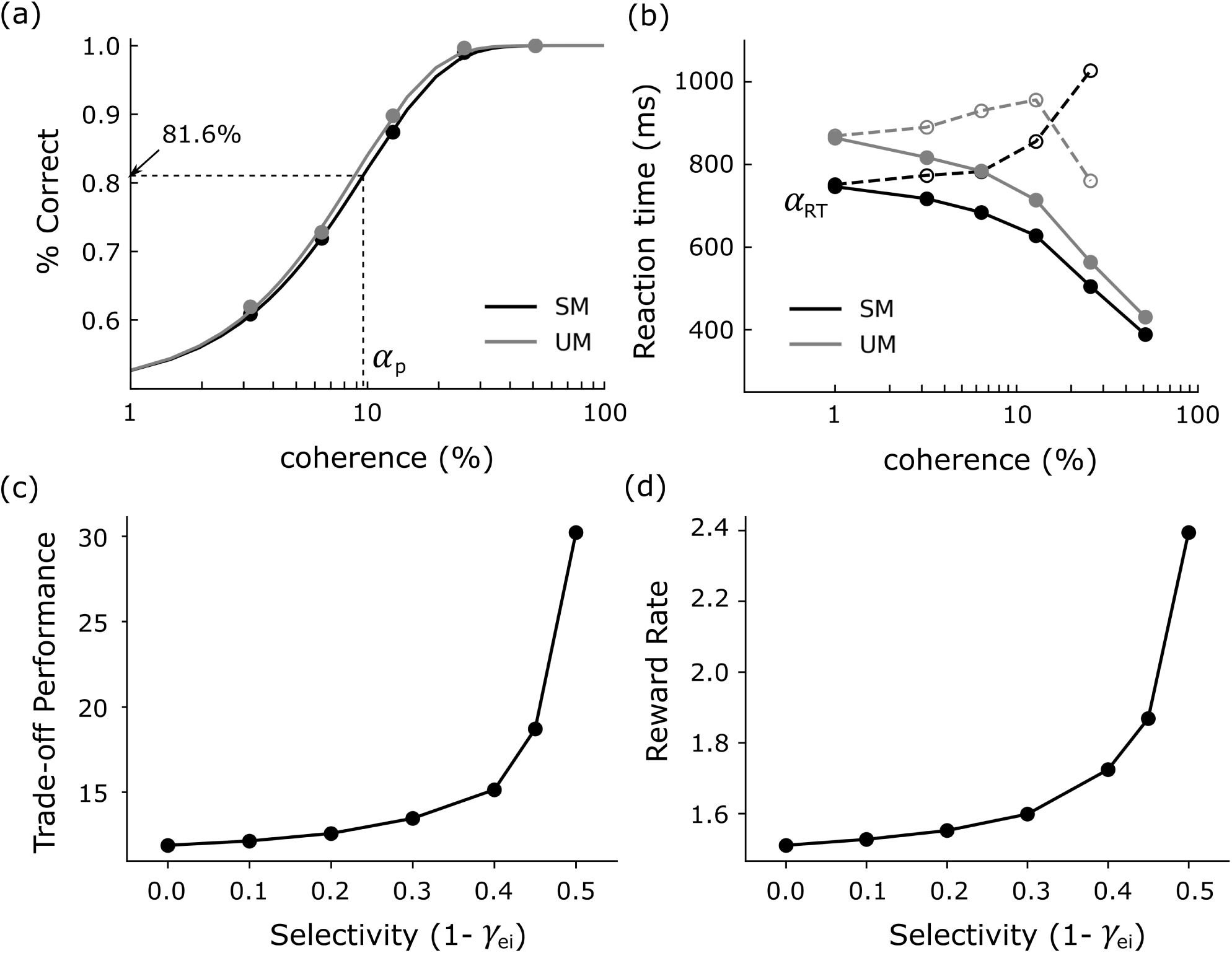
Speed-accuracy trade-off as a function of selectivity. (a) The psychometric function of a SM with (*γ*_*ee*_, *γ*_*ei*_, *γ*_*ie*_, *γ*_*ii*_) = (0.5, 0.6, 0.7, 1), alongside the psychometric function of an UM with parameters *γ*_*ee*_ = 0.5. The definition of the performance index *α*_*p*_, which is the coherence level when the model reaches 81.6% accuracy, is illustrated here. (b) The chronometric function of the SM and UM in (a). Solid circles are the reaction time for the correct trials, and hollow ones are for incorrect trials. The definition of the speed index *α*_*RT*_, which is the average reaction time at *c* = 0, is illustrated. (c) Trade-off performance, defined as the inverse of *α*_*p*_ *× α*_*RT*_, increases with selectivity, defined as 1 − *γ*_*ei*_ here. (*γ*_*ee*_, *γ*_*ie*_, *γ*_*ii*_) are fixed to be (0.5, 0.7, 1). (d) Reward rate, in units of *number of reward/s*, increases with selectivity as well. The parameters are the same as (c).

Simulation results indicate that selectivity strongly correlates with the efficiency of the trade-off (Fig. 2c, d). We observe that the trade-off performance increases along with selectivity, up until a cut-off value at *γ*_*ei*_ = 0.5 (for simplicity, in Fig. 2c, d *γ*_*ie*_ is fixed at 0.7, while only *γ*_*ei*_ is varied). This is because the selectivity of a SM cannot be unrestricted – too much selectivity leads to an inability to make decisions. Intuitively, an extreme case of *γ*_*ei*_ = *γ*_*ie*_ = 0 has no feedback inhibition from the opposing inhibitory population at all, and with insufficient mutual inhibition between inhibitory populations, there will be no “competing” mechanism that allows one population to win over the other.

### Time Varying Energy Landscape

Why does selectivity lead to a more efficient speed-accuracy trade-off? To better analyze the dynamics of our system, we first reduce the three-variable UM and the four-variable SM into a two-variable form once again, without evoking assumption (III) (see Methods). This is done by observing that the variables exhibit simple relationships after performing a coordinate transform, and these relations could be used to simplify the population models. That is, it can be shown that the dynamics of *S*_*G*_ is largely determined by the average value of *S*_*E*1_ and *S*_*E*2_, i.e. “how much” the system is activated. In fact, the relation between *S*_*G*_ and 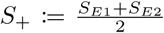 is almost linear in all time periods under many parameter sets of the UM (Fig. S3a). For the SM, the relationship between 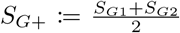 and *S*_+_ is linear, and the relation between 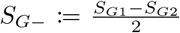and 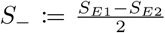 is quadratic (Fig. 3a). This means that the average inhibitory activity *S*_*G*+_ is driven by average excitatory activity *S*_+_, while the inhibitory difference *S*_*G*−_ is dependent on the excitatory difference *S*_−_.

**Figure 3:**
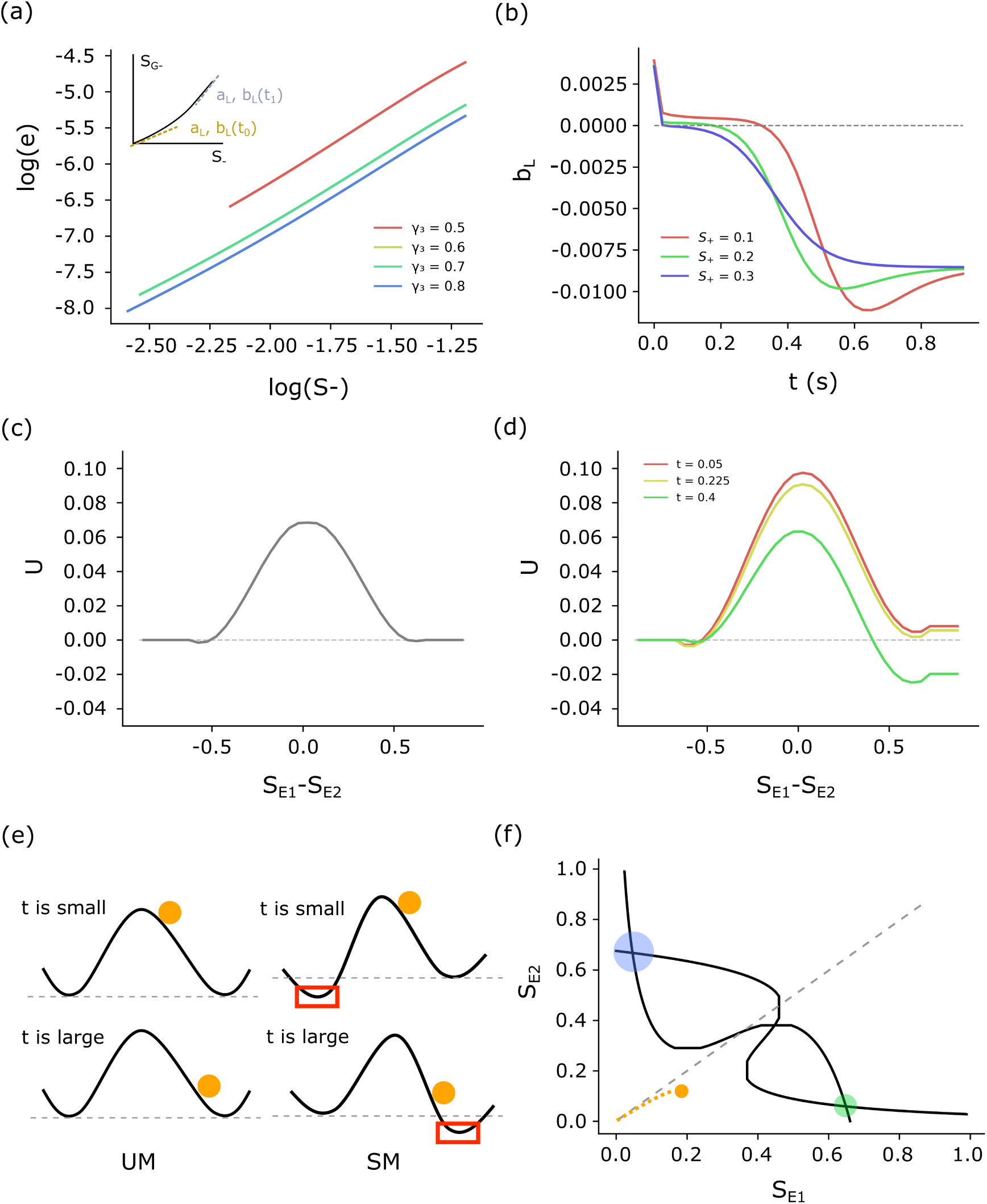
Reduced models and energy landscapes of UM and SM. (a) Quadratic relationship between *S*_−_ and *S*_*G*−_. The parameters for (*γ*_*ee*_, *γ*_*ei*_, *γ*_*ii*_) are (0.3, 0.6, 1). *log*(*e*) as a function of *log*(*S*_−_), where *e* is the residual of a linear fit between *S*_*G*−_ and *S*_−_. The slope of the resulting log-log plot is around 2, indicating that it is a quadratic function. The inset plot shows how *a*_*L*_(*t*) and *b*_*L*_(*t*) is derived from the quadratic relation. (b) Evolution of the time-varying intercept *b*_*L*_ as a function of the starting point ((*S*_*E*1_, *S*_*E*2_) = (*S*_+_ + 0.05, *S*_+_)) of the no-noise trial. In the following panels, the linearized quadratic models are indexed by the time period in which they are fitted to. (c) The energy landscape *U* of a UM with *γ*_*ee*_ = 0.5. (d) The energy landscape *U* of a SM with (*γ*_*ee*_, *γ*_*ei*_, *γ*_*ie*_, *γ*_*ii*_) = (0.5, 0.6, 0.7, 1). (e) Schematic diagram of how *U* evolves in time for the UM and SM. The orange ball indicates the position of the system. The UM landscape is fixed in time, while for the SM, the landscape favors the opposing attractor in initial stages, leading to careful weighting of options; in later stages the opposing attractor disappears, leading to faster reaction times. (f) A schematic diagram of the initial stages of the phase diagram for a SM.

Therefore, substituting *S*_*G*_ by *a*^∗^*S*_+_ + *b*^∗^ for the UM, where *a*^∗^, *b*^∗^ is the slope and intercept of the *S*_*G*_-*S*_+_ relation (Eq 6), we would again obtain a WWM. Only this time, a different synaptic weight set would result in a different reduced model. For the SM, we could similarly replace *S*_*G*−_ by 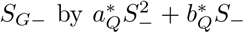, where 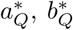 is the quadratic and linear coefficient of the *S*_*G*−_-*S*_−_ relation. A quadratic relationship can be treated as momentarily linear with a slope and intercept changing with time. Therefore, we can further reduce our SM to a linear form that resembles the UM except for a time varying slope and intercept (Eq 8). The 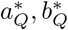 values are fitted to an unfair, noiseless trial released at (*S*_*E*1_, *S*_*E*2_) = (0.15, 0.1), and the 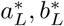 values are subsequently fitted to the data points from different time periods (see Methods). The *a*_*L*_ values slowly increase in time, while the *b*_*L*_ value is initially positive then becomes negative, as one would expect from a quadratic function with positive coefficient for the quadratic and linear terms (Fig. S5a, 3b red lines). The effect of the change in these two values can be elucidated by looking at its influence on the synaptic currents (Eq 8, restated here for convenience):

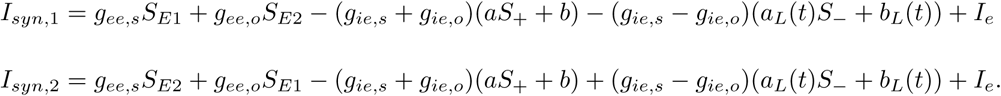

Since *g*_*ie,s*_ − *g*_*ie,o*_ is positive in the cases of interest ((*γ*_*ei*_, *γ*_*ie*_) = (0.6, 0.7)), when *a*_*L*_ increases, it slows down the system by decreasing *I*_*syn*,1_ and *I*_*syn*,2_. This is confirmed by the energy landscape of different *a*_*L*_ values, where the slope of the hill decreases for increasing *a*_*L*_ (Fig. S5b, where *b*_*L*_ is temporarily set to 0 to highlight the effect of altering *a*_*L*_ alone). Note that in contrary to the SM, the UM has a fixed energy landscape across time since it has no time varying parameters (Fig. 3c).

Unlike *a*_*L*_, which has a symmetrical effect on both of the synaptic currents, *b*_*L*_ results in an asymmetry, where positive *b*_*L*_ increases *I*_*syn*,2_ but decreases *I*_*syn*,1_. Therefore, with an initially positive *b*_*L*_, the energy landscape favors the attractor opposite of the one the system is currently attracted to. For instance, if initially *S*_*E*1_ *> S*_*E*2_, then the attractor of *S*_*E*2_ has a larger basin of attraction (Fig. 3d, red line). However, since *b*_*L*_ eventually becomes negative, this means that if the system persists in having *S*_*E*1_ *> S*_*E*2_, the tables turn and the *S*_*E*1_ attractor has a larger basin of attraction instead (Fig. 3d, green line). This change in the sizes of the basin of attractions is further discussed in Supplementary Notes S5.

Together, this means that the system carefully weighs the information in the beginning by having a larger basin of attraction at the opposing side (Fig. 3e, top right; Fig. 3f). If the evidence for the currently favored side is insufficient, the system would be attracted to the other side instead. However, if the evidence is sufficient, then the system quickly changes to a configuration that favors the current side, leading to faster convergence to the favored attractor (Fig. 3e, bottom right). The “careful consideration” part is evident in the percentage of non-regret choices, where the non-regret choices for the SM increases slower than the UM (Fig. 4a, also see Methods). That is, if *S*_*E*1_ − *S*_*E*2_ is, say, 0.04 in the UM, then it has a high chance of having *S*_*E*1_ as the winner. In contrast, for the same situation in the SM, the chance of *S*_*E*1_ winning is lower, which means that despite *S*_*E*1_ having a slight edge on *S*_*E*2_, the system is still considering its options.

**Figure 4:**
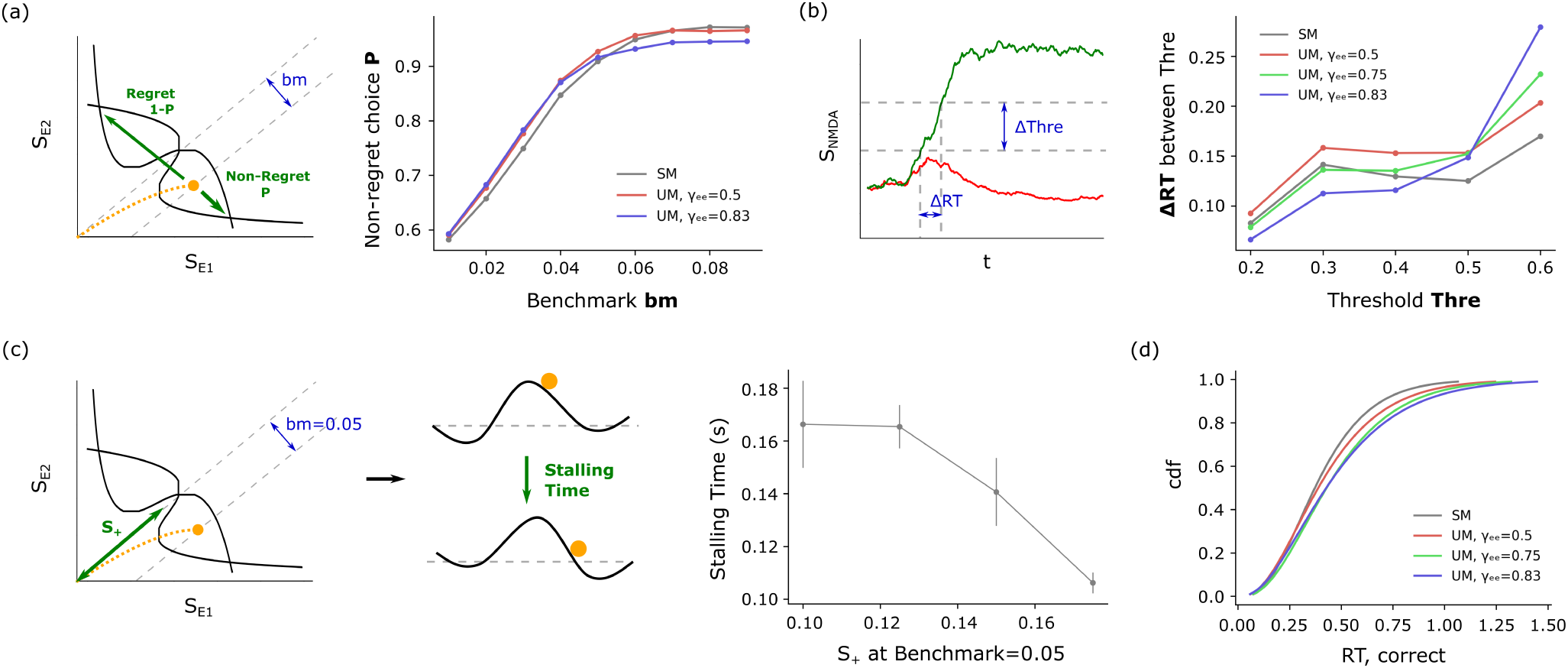
Effects of the time-varying energy landscape of SM on accuracy and reaction time. The SM for all panels uses (*γ*_*ee*_, *γ*_*ei*_, *γ*_*ie*_, *γ*_*ii*_) = (0.5, 0.6, 0.7, 1). (a) A schematic diagram shows how regret and non-regret choices are defined against a bench-mark (*bm*) (left). Percent non-regret choice as a function of benchmark (right). A lower percent non-regret choice in smaller benchmark values means that the model chooses more carefully. (b) Schematic of the average traveling time between thresholds is calculated (left). The average time it takes to reach the next threshold, starting from the current threshold (right). The x-axis is the value of the next threshold, and the starting threshold is the next threshold minus 0.1. The SM is the fastest during the later stages. (c) Schematic of how the stalling time is calculated: after hitting the benchmark, the time it takes for the energy landscape to turn from a positive *b*_*L*_ value to a negative value is the stalling time (left, middle). The smaller this value is, the faster the energy landscape turns to favor the current attractor. X-axis: *S*_+_ value of the trial when it reaches a benchmark of 0.05. Y-axis: Time until *b*_*L*_ becomes negative after the trial crosses a benchmark of 0.05 (the stalling time); mean *±* SEM (right). (d) Cumulative probability density function of the reaction time for coherence *c* = 0 trials. The distribution of the data is fit to a gamma distribution. The SM does not have a long tail, compared to the UMs.

Further statistical analysis done on the reaction time supports the conclusion of the “quick convergence” part. We first calculated the time it takes for the system to travel between thresholds (Fig. 4b). Setting the threshold to different values will result in different reaction times, and calculating the difference in reaction time between successive thresholds would allow us to know how much time it takes to reach one threshold from the previous one. Initially, the SM takes more time than some of the faster UMs to reach the next threshold. This is overturned in later stages, where the SM becomes the fastest one.

Lastly, if during the trial, the system keeps switching back and forth between *S*_*E*1_ and *S*_*E*2_, then *b*_*L*_ turns negative faster, which corresponds to strengthening the favored side faster (Fig. 3b, green and blue lines). This means that if the evidence does not favor either side, and it is taking too long to make a decision, then the system escalates the process by making it easier for the favored side to win. This can be showed by grouping trials using the *S*_+_ value at the moment it crosses a certain benchmark (set to 0.05 here to be consistent with Fig. 3b). A larger *S*_+_ value at the benchmark signifies that the trial walks the separatrix (the line where *S*_*E*1_ = *S*_*E*2_) longer than others. For each group, we can then calculate the time it takes for *b*_*L*_ to turn negative after the trial crosses the benchmark. This value decreases as *S*_+_ increases, meaning that if the trial takes too long, then *b*_*L*_ turns negative faster, which in turns means the system allows the decision to converge faster to its attractors (Fig. 4c). This could prevent the system from stalling.

The combined effect of “quick convergence” and “avoid stalling” is also evident in the distribution of the reaction time (Fig. 4d). The tail of the reaction time distribution of the SM is significantly shorter than the UMs, indicating that it has less trials with longer reaction times.

We note that the “converge quickly” portion seems to have a larger effect than the “choose carefully” portion on the speed-accuracy trade-off. That is, the changes selectivity have on the accuracy (Fig. 5a) is less notable compared to the changes in reaction time (Fig. 5b). Therefore, when evaluated in terms of the trade-off performance 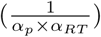, the more selective the model is, the more efficient it seems to be. However, even if we constrained the drop in accuracy to be within a certain regime, the SMs still out perform the UMs. This is evident by the fact that the SMs lie of the lower-left corner of the *α*_*p*_-*α*_*RT*_ plane, which corresponds to a more efficient speed-accuracy trade-off.

**Figure 5:**
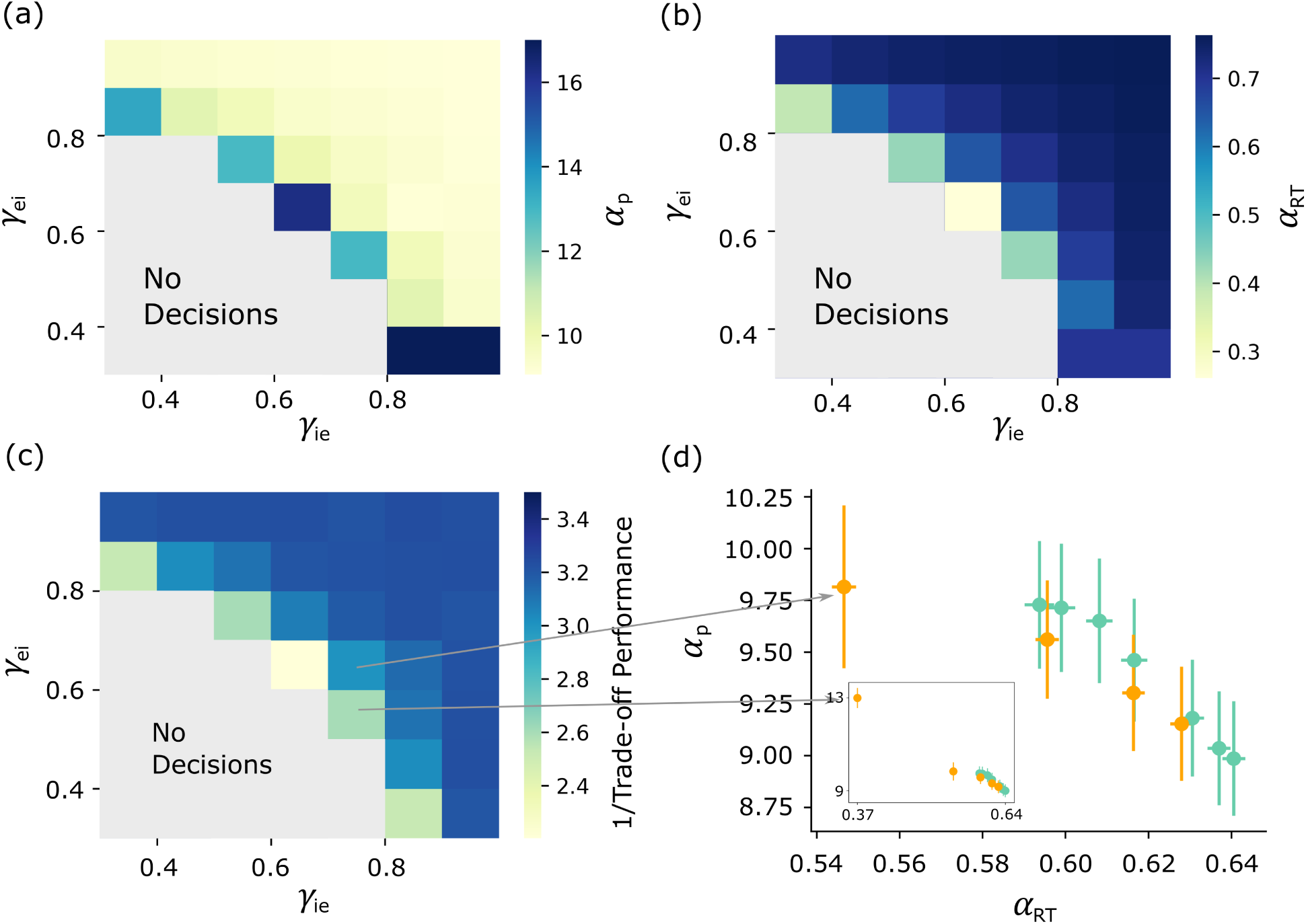
Performance index (*α*_*p*_) and speed index (*α*_*RT*_) as a function of selectivity. (a) *α*_*p*_ as a function of selectivity. (*γ*_*ee*_, *γ*_*ii*_)= (0.5, 1). Regions that cannot make decisions are labeled as “no decisions”. (b) *α*_*RT*_ as a function of selectivity. (c) Trade-off performance (the inverse of *α*_*p*_ *× α*_*RT*_) as a function of selectivity. (d) The distribution of models on the *α*_*p*_-*α*_*RT*_ plane. The green dots are UMs with different *γ*_*ee*_ values (from right to left: 0.1,0.3,0.5,0.7,0.75,0.8,0.83). The orange dots are SMs with (*γ*_*ee*_,*γ*_*ie*_,*γ*_*ii*_) and with different *γ*_*ei*_ values (from left to right: 0.6,0.7,0.8,0.9). *Inset:* The same graph with an extra data point *γ*_*ei*_ = 0.5 included.

### GABA Dynamics Underlying the Time-Varying Energy Landscape

We have shown that the key to the more efficient trade-off performance of the SM is due to its time-varying energy landscape, and that the time dependence arises from the quadratic relation of *S*_*G*−_ with *S*_−_. To better understand how the linear relation of *S*_*G*+_ with *S*_+_ and the quadratic relation of *S*_*G*−_ with *S*_−_ arises, we turn to the equations governing the GABA dynamics. According to assumption (II), since the f-I curve of the inhibitory population can be linearized, the equations of *S*_*G*1_ and *S*_*G*2_ are linear. Hence we can write down the equation of motion for *S*_*G*+_ using Eq 1 and 2:

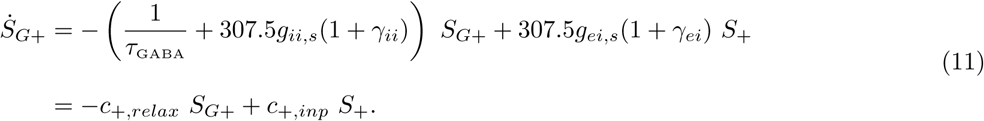

And the equation of motion for *S*_*G*−_ is:

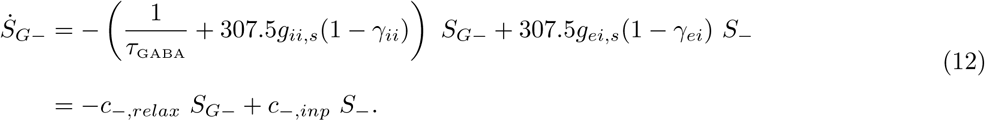

The number 307.5 comes from linearizing *ϕ*(*x*) in Eq 1 (see Eq S3). The time constants for *S*_*G*+_ and *S*_*G*−_ are 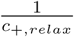 and 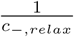 (0.001 and 0.005 in our simulations), respectively. Since *γ*_*ii*_ is always larger or equal to zero, *c*_+,*relax*_ must also be larger or equal to *c*_−,*relax*_. In the strictly larger scenario, this means that *S*_*G*+_ will always move towards steady state at a faster speed than *S*_*G*−_. This can be shown by plotting the deviation from steady state of a noiseless trial for a SM, where *S*_*G*−_ clearly deviates more than *S*_*G*+_ (Fig. 6a, bottom panel).

**Figure 6:**
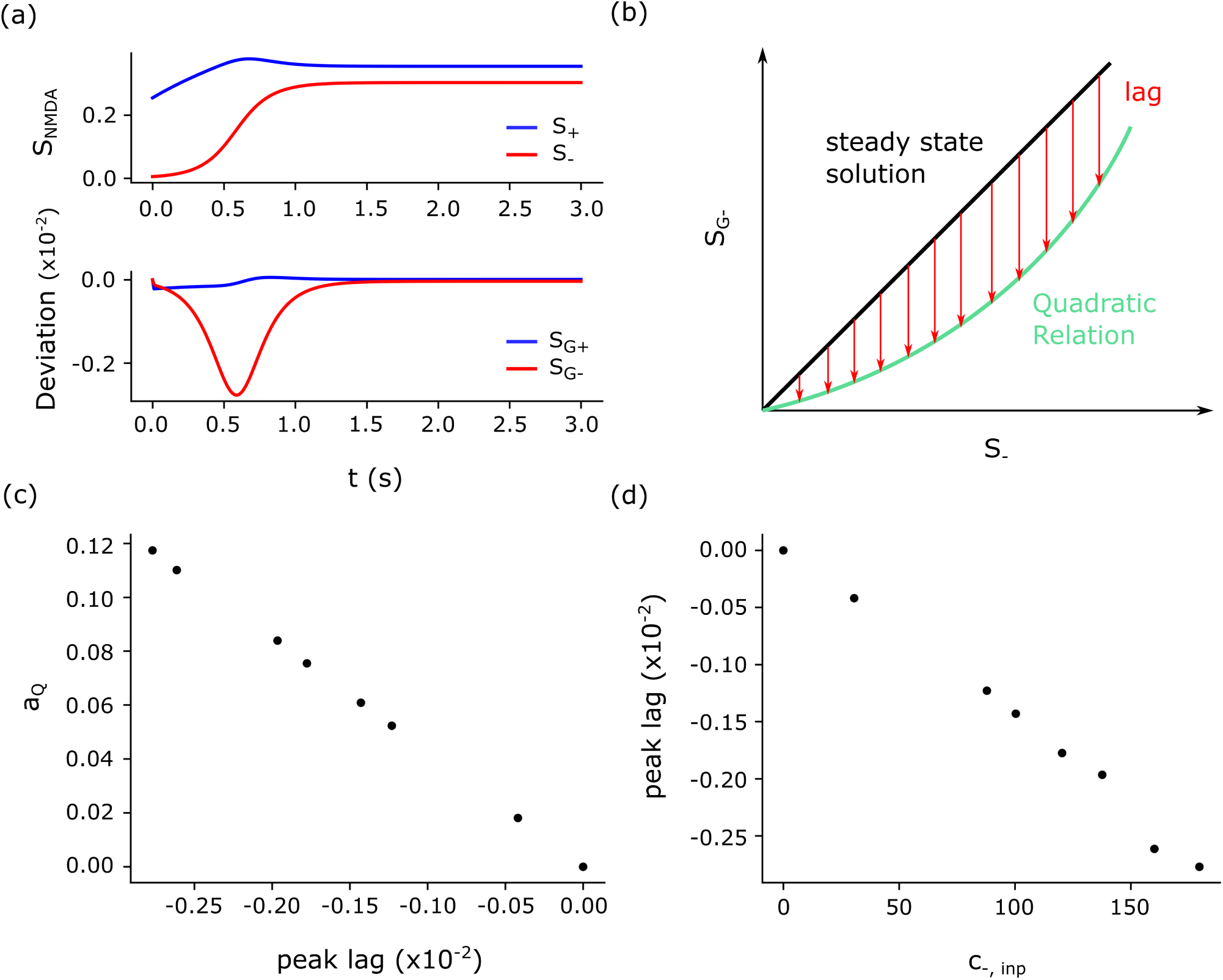
Relaxation dynamics of inhibitory average *S*_*G*+_ and inhibitory difference *S*_*G*−_. Since the inhibitory dynamics are fast, its relation with the excitatory dynamics are dictated by the evolution of *S*_+_ and *S*_−_. The lag between the inhibitory dynamics’ true state and its steady state is directly related to the “swiftness” of the relaxation term (*c*_−,*relax*_) relative to the input term (*c*_−,*inp*_). (a) *Top panel*: *S*_+_ and *S*_−_ in a biased, noiseless trial that starts at (*S*_*E*1_, *S*_*E*2_) = (0.25, 0.26). Since it is a noiseless trial, the input starts at *t* = 0 (s) instead of *t* = 0.5 (s). *Bottom panel*: The lag of *S*_*G*+_ (*S*_*G*+_ − *S*_*G*+,*ss*_, where *ss* stands for steady state) and *S*_*G*−_ (*S*_*G*−_ − *S*_*G*−,*ss*_). The evolution of *S*_+_ is linear (in the initial stages), while those of *S*_−_ is not. (b) Schematic diagram of how the quadratic relation arises from the bell-shaped lag. (c) *a*_*Q*_ as a function of the peak of the lag of *S*_*G*−_. The models, from left to right, are SMs with: (*γ*_*ee*_, *γ*_*ei*_, *γ*_*ie*_, *γ*_*ii*_) = (0.5, 0.6, 0.7, 1), (0.3, 0.6, 0.5, 1), (0.5, 0.6, 0.8, 1), (0.3, 0.6, 0.6, 1), (0.3, 0.6, 0.7, 1), (0.3, 0.6, 0.5, 1) and (0.3, 0.8, 1.25, 1). The rightmost dot is an UM with *γ*_*ee*_ = 0.5. (d) The peak of the lag as a function of *c*_−,*inp*_. The order of the models are the same as (b).

The steady state of Eq 11 and Eq 12 yields a linear relation between both *S*_*G*+_ and *S*_+_, as well as *S*_*G*−_ and *S*_−_. However, it can be shown that the deviation between the true dynamics and the steady state is non-neglectable (defined as *S*_*G*+_ − *S*_*G*+,*ss*_ for *S*_*G*+_ and *S*_*G*−_ − *S*_*G*−,*ss*_ for *S*_*G*−_, where *ss* stands for steady state; see Supplementary Notes S2 and Fig. S2e). This deviation reflects how much the true state of the system “lags” behind the steady state, and this non-trivial lag is why assumption (III) does not hold in the first place. Therefore, to gain a more accurate description for these relations, we must take into account the modifications caused by the lags. The lag of *S*_*G*+_ is roughly a linear decline, which means that *S*_*G*+_ and *S*_+_ will more or less retain its linear relation, albeit with a different slope and intercept than the steady state relation. On the other hand, the lag of *S*_*G*−_ is bell shaped, which means that the subtraction of its lag from the steady state relation will yield a concave up and increasing curve (Fig. 6b). This is the quadratic relation we observe in Fig. 3a. We can verify this by plotting *a*_*Q*_ as a function of the peak of the lag (Fig. 6c).

The shape of the lags are in turn a result of the velocity flow of *S*_+_ and *S*_−_ (Fig. 6a, top panel). While the velocity of *S*_+_ is roughly linear with a slight decline late in the trial, the velocity of *S*_−_ is hyperbolic tangent like. This means that as the speed of *S*_−_ increases in the initial stages, it becomes more challenging for *S*_*G*−_ to keep up, resulting in an increasing lag. As the speed decreases, the lag then becomes less severe. This would thus result in a bell shaped lag. Therefore, to obtain a more quadratic curve, the peak of the lag must be large. The magnitude of the lag is controlled by both *c*_−,*relax*_ and *c*_−,*inp*_. In our models, *g*_*ii,s*_ and *γ*_*ii*_ are fixed, so every model has the same *c*_−,*relax*_ value. In that case, the lag will be proportional to *c*_−,*inp*_, as we indeed see in Fig. 6d.

## Discussion

We proposed a more generalized model (SM) for decision making that reproduced behavioral and neuro-physiological observations. We discovered that selectivity in the inhibitory population creates a time-dependent energy function, which allows the model to reach a more efficient speed-accuracy trade-off. This includes weighing the choices more carefully in initial stages, converging to an attractor quickly once the decision is made, and avoiding long reaction time trials. Furthermore, it emphasizes the importance of the role the GABA dynamics play, despite being 20 times faster than NMDA dynamics. This topic can be further extended in several aspects. First, the SM introduces a rudimentary level of generalization of the attractor model. Specifically, it is found that heterogeneity is commonly observed in brain areas associated with decision making (Rigotti et al., 2013; Jun et al., 2010), and sub-dividing the inhibitory populations into two is a first step towards the inclusion of heterogeneity. We showed that generalizing the model by having heterogeneous synapses improves the decision making ability of the network. If this one step of separation elicits different dynamics, we could expect more complicated forms of functionality from an even higher level of heterogeneity. Together with other studies that demonstrates how heterogeneity aids in encoding information regarding different aspects of the stimulus (Aoi et al., 2020), this highlights the importance of incorporating heterogeneity into future decision making models.

Another important concept from decision making literature is excitation-inhibition balance. E/I balance could refer to either balanced top-down input currents, or the balance of excitation-to-inhibition synaptic currents for the neurons. In terms of the former, it is previously shown that a network with balanced inputs is capable of making more accurate decisions via having a time-varying energy landscape (Wang et al., 2013). This time-variance is different from those mentioned in our study, as the variations of the landscape occurs symmetrically to both of the attractors in this case, and asymmetrically in our case. Furthermore, this time-variance requires external top-down control (Wang et al., 2013; Lo et al., 2015), while in the SM, this property is naturally embedded in local circuits whose configuration is supported by the latest experimental observations. The interaction between SM and balanced external inputs, both of which are time-varying components on their own, requires further investigation.

The balancing of the synaptic currents too warrant a deeper look. In our simulations, altering synaptic weights result in drastically different dynamics, as can be seen from the results of altering *γ*_*ee*_. In fact, altering a single synaptic weight by 0.5% can significantly speed up or slow down the reaction time of the system. Previous studies have shown that an imbalance in excitation/inhibition ratio changes the system’s behavior: over-excited, and the system tends towards impulsive decisions; over-inhibited, and it becomes indecisive (Lam et al., 2017). This picture of balanced excitation/inhibition ratio is a widely explored topic in the field of neuroscience, both experimentally and theoretically. The introduction of two inhibitory populations adds another layer of complexity to the balanced picture, as the balance is now controlled by two inhibitory populations and their interactions simultaneously. How E/I balance may come into play after this complication, and how the dynamics may change accordingly, is an open question.

The process of learning how to perform these decision tasks is another topic that has gained attention (Roy et al., 2021). An interesting result in the study of Najafi *et al*. (2020) is that equal selectivity between excitatory and inhibitory populations is maintained throughout the course of learning. Is selectivity a property that manifests naturally as a consequence of learning? Specifically, given that SMs are more efficient in their speed-accuracy trade-off than the UMs, could selectivity emerge out of a non-selective network automatically as it learns? Could the network self-organize into a more efficient state of selectivity?

Aside from broad concepts of heterogeneity, E/I balance and learning, another piece of literature to integrate or compare the SM to would be other types of decision making models. In particular, we note how the property of the time-varying landscape has an “urgency” component for trials that take a longer time. That is, if the system is indecisive for too long, the landscape would adjust itself so that convergence to an attractor becomes easier. One of the decision making models proposed in the past is the urgency gating model, which shares the same concept of having an urgency signal that rushes the decision making process along in later stages. However, the SM is mechanistically different from the urgency gating model, which bases decisions on momentary information instead of integrated information (Ditterich, 2006; Cisek et al., 2009; Thura et al., 2012). In this regard, the SM is more similar to the diffusion model, with differences such as time-shift invariance pointed out in previous studies (Wong et al., 2007; Wong and Huk, 2008). A more thorough analysis of experimental evidence for each of these models may shed light on how these models may be related to one another.

Last but not least, the details that are uncovered in this study are also worth pursuing. For instance, the new reduction reduces the UM and SM back to two variables in a more realistic way. How might the linear or quadratic relationship depend on the synaptic weights? Why do such small deviations from linearity impact the dynamics? How does the GABA dimension change the energy landscape of the system? In our studies, the contribution of AMPA is neglected. However, our results suggest that the inclusion of GABA dynamics significantly alter the decision making network’s accuracy and reaction time; hence it is entirely probable that an inclusion of AMPA would also impact the dynamics. In conclusion, our study supports the idea of a more complicated picture of decision making models than the widely studied UM, an adjustment that leads to better computational abilities.

## Supporting information

attractor_selectivity_supp

## Acknowledgements

We thank Alexander J. White, Cheng-Te Wang and Ching-Che Charng for discussion and feedback on the manuscript. This work was partially supported by Ministry of Science and Technology grant no. 110-2218-E-007-035 and grant no. MOST 110-2112-M-007-003-, and by the Featured Areas Research Center Program within the framework of the Higher Education Sprout Project, a join fund from the Ministry of Education and the Ministry of Science and Technology in Taiwan. The authors also gratefully acknowledge the support from the National Center for Theoretical Sciences, Taiwan.

