## Supplementary material for "Choose carefully, act quickly: Efficient decision making with selective inhibition in attractor neural networks": attractor_selectivity_supp

### Supplementary Notes

#### S1 WWM (Wong and Wang's Model)

The derivation of the WWM is given in Wong and Wang (2006), and here we only summarized the results. The WWM takes the following form:

$$\begin{aligned}\dot{S}_{E1} &= -\frac{S_{E1}}{\tau_{\text{NMDA}}} + 0.641 \times (1 - S_{E1})H(I_{syn,1}) \\ \dot{S}_{E2} &= -\frac{S_{E2}}{\tau_{\text{NMDA}}} + 0.641 \times (1 - S_{E2})H(I_{syn,2}).\end{aligned} \tag{S1}$$

$H(x)$ can be found in Wong and Wang (2006), but is restated here for convenience:

$$H(x) = \frac{270x - 108}{1 - e^{-0.154\times(270x-108)}}. \tag{S2}$$

The f-I curve for the inhibitory population, $\phi(x)$ Wong and Wang (2006), is not used in the WWM but is used in the UM and SM Wong and Wang (2006):

$$\phi(x) = 307.5x - 88.5. \tag{S3}$$

The synaptic current takes the form

$$\begin{aligned}I_{syn,1} &= J_e S_{E1} - J_i S_{E2} + I_b + I_n + I_1 \\ I_{syn,2} &= J_e S_{E2} - J_i S_{E1} + I_b + I_n + I_2,\end{aligned} \tag{S4}$$

where $J_e$ is the self-exciting synaptic weight, $J_i$ is the inhibitory synaptic weight between the two populations, $I_b$ is the bias current of the two populations, and $I_1$, $I_2$ represents the input signal (see Fig. 1a). $I_n$ is an Ornstein-Uhlenbeck process, which is a Gaussian white noise signal with its higher frequencies attenuated. It takes the form:

$$\dot{I}_n = -I_n + \mathcal{N}(0,1) \times \frac{\sqrt{D^2\Delta}}{\Delta}, \tag{S5}$$

where $\Delta = 0.002$ (which matches the time constant of AMPA), $D = 5$ (in our paper), and $\mathcal{N}(0,1)$ represents a standard Gaussian.

#### S2 Non-trivial GABA Dynamics

The WWM assumes $\frac{dS_G}{dt} = 0$ (assumption (III)), which can be shown not hold in general cases. This is done by constructing different UMs (SMs works as well, but is unnecessary for this proof of concept) that collapse back into the same WWM upon activating assumption (III). Here we first describe how these UMs are constructed, then discuss the simulation results using these UMs.

Assuming assumption (III) to be true, we can reduce the differential equation of $S_G$ into an algebraic equation, and combine it with the differential equations for the NMDA channels. Since $I_{syn,1}$ is a linear combination of $S_{E1}$, $S_{E2}$ and $S_G$, and $S_G$ is a linear combination of $S_{E1}$ and $S_{E2}$, the substitution would result in a linear combination of $S_{E1}$, $S_{E2}$ and a bias term (see Methods). Therefore, we could collect the terms for the coefficients of $S_{E1}$, $S_{E2}$ and the bias, and map it onto $\{J_e, J_i, I_b\}$. Enforcing constraints (I)-(IV) to ensure the mapping is one-to-one (see Methods), we can obtain unique UM parameters by solving the following equations:

$$\begin{aligned} g_{ee,s} - \frac{1.5375 g_{ie}^2}{1 + 1.5375 g_{ii}} &= J_e \\ \gamma_{ee} g_{ee,s} - \frac{1.5375 g_{ie}^2}{1 + 1.5375 g_{ii}} &= -J_i \\ I_e - \frac{1.1525 g_{ie}}{1 + 1.5375 g_{ii}} &= I_b. \end{aligned} \tag{S6}$$

Similarly, we can obtain unique SM parameters by solving:

$$\begin{aligned} g_{ee,s} - \frac{g_{ie,s}}{d}\left(1.5375 g_{ei,s} - \frac{1.5375^2 g_{ii,o} g_{ei,o}}{1 + 1.5375 g_{ii,s}}\right) - \frac{g_{ie,o}}{d}\left(1.5375 g_{ei,o} \frac{1.5375^2 g_{ii,o} g_{ei,o}}{1 + 1.5375 * g_{ii,s}}\right) &= J_e \\ g_{ee,o} - \frac{g_{ie,s}}{d}\left(1.5375 g_{ei,o} \frac{1.5375^2 g_{ii,o} g_{ei,o}}{1 + 1.5375 * g_{ii,s}}\right) - \frac{g_{ie,o}}{d}\left(1.5375 g_{ei,s} - \frac{1.5375^2 g_{ii,o} g_{ei,o}}{1 + 1.5375 g_{ii,s}}\right) &= J_i \\ \frac{-1}{d}\left(1 - \frac{1.5375 g_{ii,o}}{1 + 1.5375 g_{ii,s}} \left(1.1525 + 1.5375 I_i\right)\left(g_{ie,s} + g_{ie,o}\right) + I_e\right) &= I_b, \end{aligned} \tag{S7}$$

where $d = 1 + 1.5375 g_{ii,s} - \frac{(a g_{ii,o})^2}{1 + a g_{ii,s}}$.

After devising a mapping that allowed us to adjust one parameter $\gamma_{ee}$ and have the other parameters take values that would allow the network to reduce to the WWM upon activating assumption (III), we then calculated the accuracy and reaction times of the UMs with different $\gamma_{ee}$ values (Fig. S2a-c). Here, the number of trials as well as the noise sequence given are identical across all UMs, thereby ruling out the uncertainty in results that might be elicited by randomly assigning noise sequences.

We showed that both the accuracy and reaction time of the different UMs varied in a non-trivial scale as $\gamma_{ee}$ is adjusted. In particular, the reaction time of incorrect trials can vary up to 0.1 (s) (Fig. S2c). To verify that this difference is indeed caused by a violation of assumption (III), we increased the speed of the GABA dynamics ten-fold:

$$\dot{S}_G = [-\frac{S_G}{\tau_{GABA}} + \phi(I_{syn,G})] \times 10. \tag{S8}$$

As expected, the accuracy and decision time converged to the original WWM (Fig. S2d). This means that although the NMDA time constant is 20 times slower than the GABA time constant, the GABA dynamics still cannot be assumed to reach steady state instantaneously. In fact, for any given time $t$ in the simulation, we can calculate how much $S_G$ lags behind its steady state value $S_G^*$. $S_G^*$ can be calculated by substituting the $S_{E1}(t)$ and $S_{E2}(t)$ values at the given time point into Eq 1, and letting $\dot{S_G} = 0$. The lag is then defined as $S_G - S_G^*$.

The lag as a function of time for different $\gamma_{ee}$ values can then be examined (Fig. S2e). We can see that the lag is small compared to the value of simulated $S_G$ value (around $\mathcal{O}(\frac{1}{100})$), as expected since it is 20 times the speed of NMDA. Nevertheless, it has a huge impact on the network's dynamics, as reflected on the accuracy and reaction time, as well as its velocity as a function of time (Fig. S2f). The magnitude of the velocity is simply the distance the system has moved from one time point to the other, divided by the time difference. As $\gamma_{ee}$ increases, the network lags further from its steady state, making its velocity larger and its decision more impulsive.

Therefore, as the GABA dimension cannot be neglected, we can expect the UM and SM to have different dynamics.

#### S3 Reduced Models

Here we determine whether the reduced models are reflective of the full model or not. We start with the linear reduced model of the UM. We gave the trials in the linear reduced and full models matching noise sequence, and compared its resulting accuracy and reaction time. Essentially, this asks whether the full and linear reduced model converge to the same choice using the same amount of time, given the same noise sequence. For some parameter sets, its reaction time across most trials are quite similar (Fig. S3f). As the $\gamma_{ee}$ value increases, the linear reduced model becomes less accurate in terms of individual trials. However, its averaged reaction time still approximately matches the full model. However, when comparing the linear reduced model of the SM to its full model, we note that the reaction time is systematically overestimated across almost all trials (data not shown).

The reason the linear reduced models predicted the SMs poorly is because in those cases, the quadratic relation between $S_{G-}$ and $S_-$ cannot be approximated by a linear relation. In fact, when fitting them to a time-dependent linear reduced model, the new reduced model predicts the accuracy and reaction time of the full model well (Fig. S3g). The mean reaction time of these models are summarized in Fig. S3h.

This states a crucial finding – the UMs (and some of the SMs whose quadratic coefficients are smaller) are well approximated by assuming a linear relation between $S_{G-}$ and $S_-$ (a first-order approximation), while others require quadratic (or second order) relations.

#### S4 Linear Reduced Models

Here we look at the speed accuracy trade-off for models that may be linearly reduced. Since the $\{J_{e,lin}, J_{i,lin}, I_{b,lin}\}$ values of the linear reduced model may not necessarily coincide with the parameters of the WWM $\{J_e, J_i, I_b\}$, it is possible that there may be some $\{J_{e,lin}, J_{i,lin}, I_{b,lin}\}$ values that are reachable to the SM but not to the UM. This turns out to not be the case: as long as the $\gamma_{ee}$ value is set to a sufficiently small value, the parameters of the WWM and linear reduced model will be close. For instance, using $\gamma_{ee} = 0.1$, the differences between the parameters predicted by the linear reduced model and the original model are $(\Delta J_e, \Delta J_i, \Delta I_b) = (-0.0002, 0.0002, 0.0002)$ (where $\Delta J_e = J_{e,lin} - J_e$, $\Delta J_i = J_{i,lin} - J_i$ and $\Delta I_b = I_{b,lin} - I_b$). Perturbing the parameters on this order of magnitude elicits only a small change in the accuracy and reaction times (Table S1). This result implies that for the parameter sets that are reducible to linear reduced models, there is not much computational differences between the UM and the SM.

#### S5 Psychometric Functions of the Time-Dependent Linear Reduced Model

Initially, the time-dependent linear reduced model has a larger basin of attraction on the opposing side. In later stages, this flips and the opposing basin of attraction shrinks. This is shown not only in the energy landscape, but in the psychometric functions of the time-dependent reduced model as well. We can create multiple linear reduced models (that are time independent) by taking $\{a_L(t), b_L(t)\}$ values at different time points, and plot their psychometric functions. For models with $\{a_L(t), b_L(t)\}$ values where $t$ is small, they have accuracy less than 0.5 at coherence $c = 0$, which corresponds to the system favoring the other side (Fig. S5d, red). Models with $\{a_L(t), b_L(t)\}$ values where $t$ is large have accuracies of 1 for all coherence levels, which signifies a disappearance of the attractor of the less favored side (Fig. S5c,d, purple).

### Supplementary Tables

| Differences in $\alpha_p$ and $\alpha_{RT}$ upon perturbing parameters | | |
|---|---|---|
| Perturbation | $\Delta\alpha_p$ | $\Delta\alpha_{RT}$ |
| $J_e + 0.0001$ | 0.0736 | 0.0008 |
| $J_i + 0.0001$ | 0.0200 | 0.0006 |
| $I_b + 0.0001$ | 0.1720 | 0.0035 |

Table S1: Changes in accuracy ($\Delta\alpha_p$) and reaction time ($\Delta\alpha_{RT}$) when perturbing $\{J_e, J_i, I_b\}$. In this order of magnitude of perturbation, the change in both $\Delta\alpha_p$ and $\Delta\alpha_{RT}$ are small, which means that there is little difference between the WWM and the UM/SMs.

### Supplementary Figures

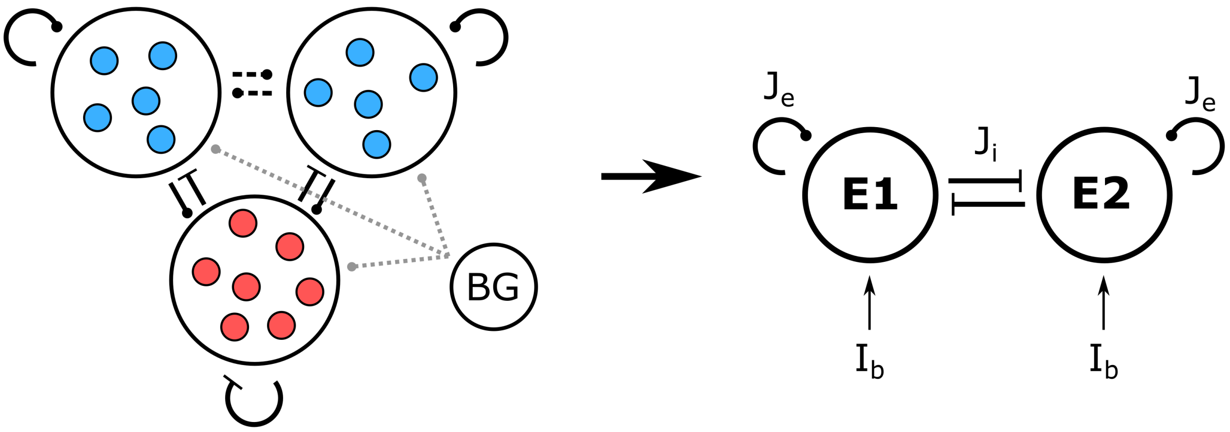


Figure S1: Reduction from a large spiking network to a WWM. BG represents the background population (those not involved with the decision making process). *Left:* spiking network model; *Right:* WWM.

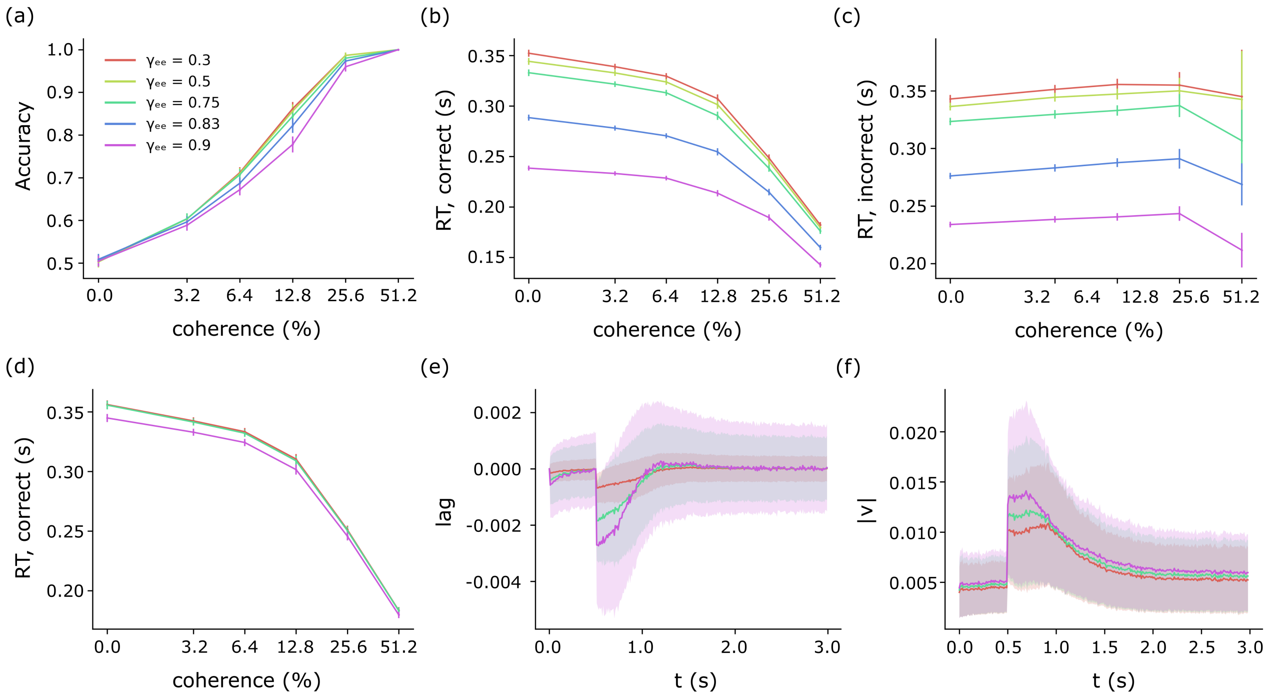


Figure S2: Accuracy, reaction time, lag and velocity of different UMs. As $\gamma_{ee}$ increases, the accuracy and reaction time both decline, while the lag and velocity increases. (a) Accuracy as a function of coherence for UMs with different $\gamma_{ee}$ values; mean $\pm$ SEM. The color scheme is consistent throughout the panels of this figure. (b) Reaction time $RT$ (of correct trials) as a function of coherence; mean $\pm$ SEM. (c) Reaction time $RT$ (of incorrect trials) as a function of coherence; mean $\pm$ SEM. (d) Reaction time $RT$ (of correct trials) as a function of coherence when the GABA dynamics are ten times faster; mean $\pm$ SEM. Under this condition, the different models have almost identical reaction times. (e) The lag of the system as a function of time; mean $\pm$ SD. (f) Magnitude of velocity of the system as a function of time; mean $\pm$ SD.

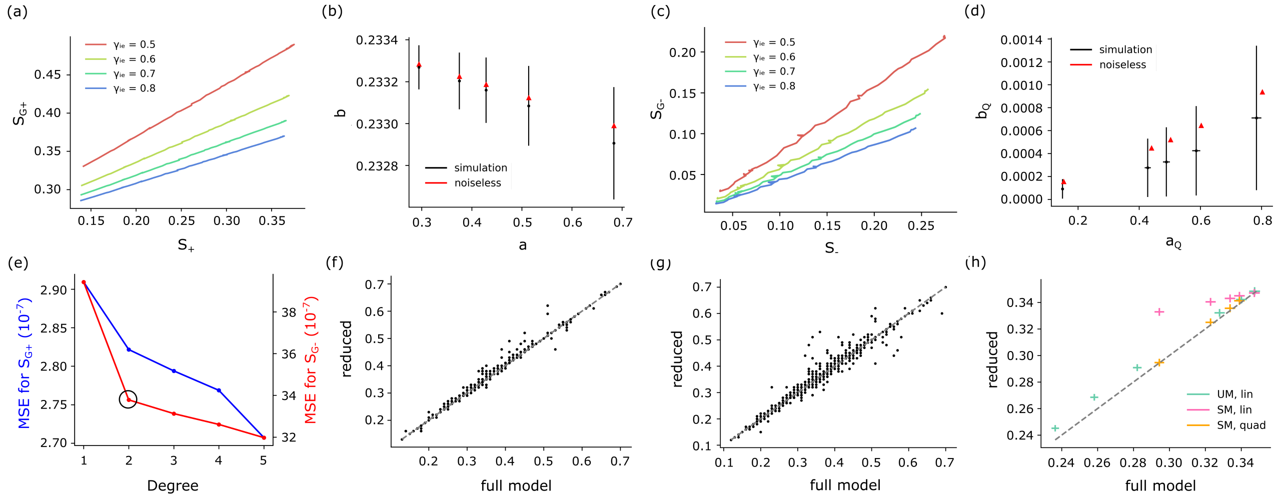


Figure S3: Empirical relations of the simulated SM, and the comparison of the full model with the reduced models, using the same noise sequences. The relation between excitatory average ($S_+$) and inhibitory average ($S_{G+}$) are shown to be linear, and the relation of excitatory difference ($S_-$) and inhibitory difference ($S_{G-}$) is quadratic. (a) Linear relationship between $S_+$ and $S_{G+}$. The parameters for $(\gamma_{ee}, \gamma_{ei}, \gamma_{ii})$ are $(0.3, 0.6, 1)$. (b) Slope ($a^*$) and intercept ($b^*$) for the linear relation during noiseless trials, compared to the slope ($a_{sim}$) and intercept ($b_{sim}$) of trials with noise; mean ± SD. The parameters for $(\gamma_{ee}, \gamma_{ii})$ are $(0.3, 1)$, and the $(\gamma_{ei}, \gamma_{ie})$ values from left to right are $(0.8, 1.25)$, $(0.6, 0.8)$, $(0.6, 0.7)$, $(0.6, 0.6)$, $(0.6, 0.5)$. (c) Quadratic relationship between $S_-$ and $S_{G-}$. The parameters are the same as (a). (d) Quadratic ($a^*_Q$) and linear coefficient ($b^*_Q$) for the linear relation during noiseless trials, compared to the quadratic ($a_{Q,sim}$) and linear coefficients ($b_{Q,sim}$) of trials with noise; mean ± SD. The order of the parameters are the same as (b). (e) Mean squared error (MSE) of $S_{G+}$ and $S_{G-}$ when fitted to different degrees of polynomials. The SM used here has $(\gamma_{ee}, \gamma_{ei}, \gamma_{ie}, \gamma_{ii}) = (0.5, 0.6, 0.7, 1)$. Using the elbow method, we know that the best fit for the $S_{G+}$-$S_+$ relation is linear (because there is no significant drop in MSE as the degree is varied), while the best fit for the $S_{G-}$-$S_-$ relation is quadratic (significant drop as degree increases becomes 2). The error bars are omitted for ease of visualization. (f) Comparison of reaction time between full and linear reduced UM model with parameter $\gamma_{ee} = 0.3$. (g) Comparison of reaction time between full and quadratic reduced SM model with parameter $\gamma_{ee} = 0.3$, $\gamma_{ei} = 0.6$, $\gamma_{ie} = 0.7$, $\gamma_{ii} = 1$. (h) Comparison of average reaction time between full and linear/quadratic reduced UM/SM models with different parameter sets. The green dots are UMs, the pink are SMs with linear reduced models, and the orange are SMs with time-dependent linear reduced models. The parameters for the UM from right-most to left are: $\gamma_{ee} = 0.1, 0.3, 0.5, 0.75, 0.8, 0.83$. The parameters for the SM from right to left are: $(\gamma_{ei}, \gamma_{ie}) = (0.8, 1.25)$, $(0.6, 0.8)$, $(0.6, 0.7)$, $(0.6, 0.6)$, $(0.6, 0.5)$; mean ± SEM.

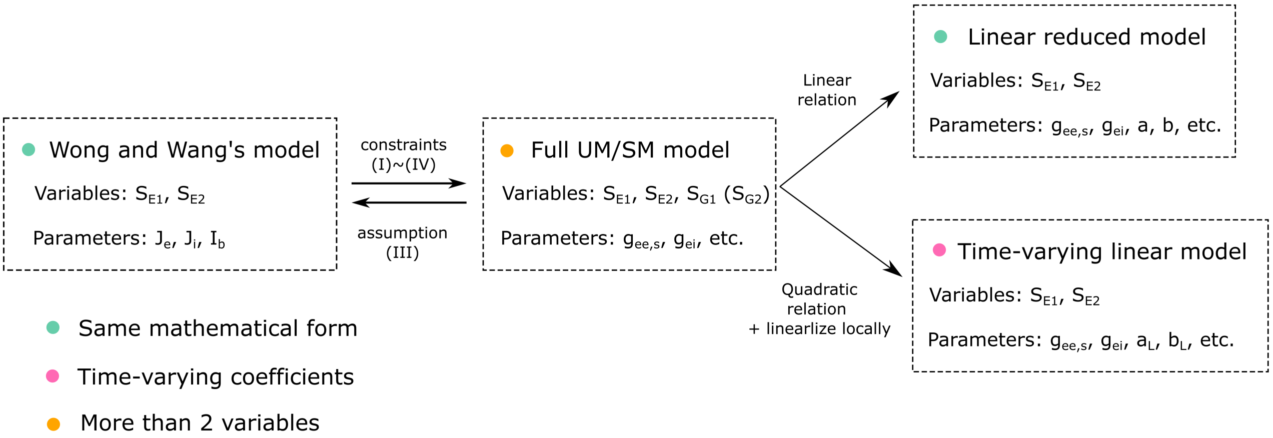


Figure S4: Illustration of the relation between the WWM, full UM/SM models and the reduced models. The full UM/SMs can be reduced to the WWM through assumption (III), and reduced to the linear and time dependent linear reduced models using linear or quadratic empirical relations between excitatory and inhibitory difference.

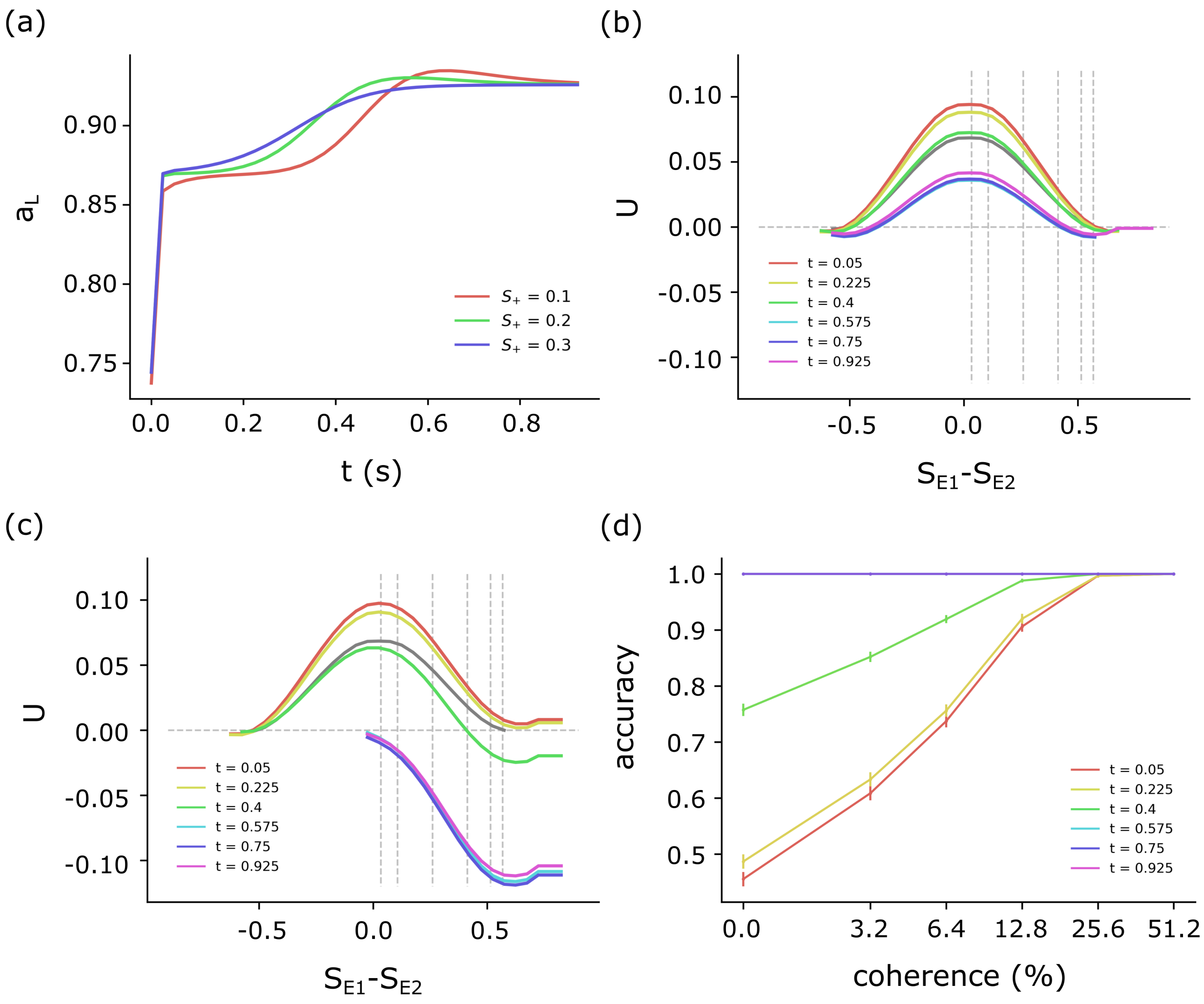


Figure S5: Time-varying energy landscape of the SM. (a) Time-varying slope $a_L$ as a function of the starting point ($(S_{E1}, S_{E2}) = (S_+ + 0.05, S_+)$) of a no-noise trial. (b) The energy landscape $U$ of a SM with $(\gamma_{ee}, \gamma_{ei}, \gamma_{ie}, \gamma_{ii}) = (0.5, 0.6, 0.7, 1)$, but with $b_L = 0$ to isolate the effect $a_L$ has on the landscape. (c) The energy landscape $U$ of a SM with $(\gamma_{ee}, \gamma_{ei}, \gamma_{ie}, \gamma_{ii}) = (0.5, 0.6, 0.7, 1)$, without setting $b_L$ to zero. (d) The psychometric function of different $\{a_L, b_L\}$ values; mean $\pm$ SEM.
